# The iron-dependent repressor YtgR regulates the tryptophan salvage pathway through a bipartite mechanism of transcriptional control in *Chlamydia trachomatis*

**DOI:** 10.1101/322586

**Authors:** Nick D. Pokorzynski, Amanda J. Brinkworth, Rey A. Carabeo

**Author notes:** Corresponding Author: Rey A. Carabeo 1770 NE Stadium Way, Biotechnology & Life Sciences Bldg., Washington State University Pullman, WA, 99164.

## Abstract

During infection, pathogens are starved of essential nutrients such as iron and tryptophan by host immune effectors. Without conserved global stress response regulators, how the obligate intracellular bacterium *Chlamydia trachomatis* arrives at a physiologically similar “persistent” state in response to starvation of either nutrient remains unclear. Here, we report on the iron-dependent regulation of the *trpRBA* tryptophan salvage pathway in *C. trachomatis*. Iron starvation specifically induces *trpBA* expression from a novel promoter element within an intergenic region flanked by *trpR* and *trpB.* YtgR, the only known iron-dependent regulator in *Chlamydia,* can bind to the *trpRBA* intergenic region upstream of the alternative *trpBA* promoter to repress transcription. Simultaneously, YtgR binding promotes the termination of transcripts from the primary promoter upstream of *trpR.* This is the first description of an iron-dependent mechanism regulating prokaryotic tryptophan biosynthesis that may indicate the existence of novel approaches to gene regulation and stress response in *Chlamydia.*

## Introduction

Nutrient acquisition is critical for the success of pathogenic bacteria. Many pathogenic bacteria must siphon nutrients from their hosts, such as nucleotides, amino acids and biometals (Brown, Palmer, & Whiteley, 2008; Eisenreich, Dandekar, Heesemann, & Goebel, 2010; Ray, Marteyn, Sansonetti, & Tang, 2009; Skaar, 2010). This common feature among pathogens renders them susceptible to nutrient limitation strategies associated with the host immune response (Hood & Skaar, 2012). Counteractively, bacterial pathogens have evolved sophisticated molecular mechanisms to respond to nutrient deprivation, involving increasingly complex and sophisticated nutrient-sensing regulatory networks. These stress response mechanisms are essential for pathogens to avoid clearance by the immune system. By delineating their function at the molecular level, we can better target aspects of the host-pathogen interface suitable for therapeutic manipulation. However, stress responses in the obligate intracellular bacterium *Chlamydia trachomatis* are relatively poorly characterized, leaving unanswered many fundamental questions about the biology of this pathogen.

*Chlamydia trachomatis* is the leading cause of bacterial sexually transmitted infections (STIs) and infection-derived preventable blindness worldwide (CDC, 2017; Newman et al., 2015; H. R. Taylor, Burton, Haddad, West, & Wright, 2014). Genital infections of chlamydia disproportionately affect women and are associated with serious sequelae in the female reproductive tract such as tubal factor infertility (Hafner, 2015). Chlamydiae are Gram-negative bacterial parasites that develop within a pathogen-specified membrane-bound organelle termed the inclusion (Moore & Ouellette, 2014). Chlamydial development is uniquely characterized by a biphasic interconversion of an infectious elementary body (EB) with a non-infectious, but replicative reticulate body (RB) (AbdelRahman & Belland, 2005). An obligate intracellular lifestyle has led to reductive genome evolution across chlamydial species; Chlamydiae have retained genes uniquely required for their survival, but have become nutritionally dependent on their hosts by discarding many metabolism-related genes (Clarke, 2011). Of note, *C. trachomatis* does not possess genes necessary for eliciting a stringent response to nutrient starvation (*e.g. relA, spoT*), suggesting that this pathogen may utilize novel mechanisms to respond to nutrient stress (Stephens et al., 1998).

It is well established that in response to various stressors, Chlamydiae deviate from their normal developmental program to initiate an aberrant developmental state, termed “persistence” (Wyrick, 2010). This persistent state is distinguished by the presence of viable, but non-cultivable, abnormally enlarged chlamydial organisms that display dysregulated gene expression. Importantly, *Chlamydia* can be reactivated from persistence by abatement of the stress condition. As such, chlamydial persistence at least superficially resembles a global stress response mechanism. Yet the molecular underpinnings of this phenotype are poorly understood, with most published studies focusing on the molecular and metabolic character of the aberrant, persistent form. It is therefore unclear to what extent primary stress responses contribute to the global persistent phenotype in *Chlamydia.*

The best described inducer of persistence is the pro-inflammatory cytokine interferon-gamma (IFN-γ). The bacteriostatic effect of IFN- γ has been primarily attributed to host cell tryptophan (Trp) catabolism, an amino acid for which *C. trachomatis* is auxotrophic (Byrne, Lehmann, & Landry, 1986; Fehlner-Gardiner et al., 2002; M. W. Taylor & Feng, 1991). Following IFN- γ stimulation, infected host cells up- regulate expression of indoleamine-2,3-dioxygenase (IDO1), which catabolizes Trp to W-formylkynurenine via cleavage of the indole ring (Macchiarulo, Camaioni, Nuti, & Pellicciari, 2009). *C. trachomatis* cannot recycle kynurenines, unlike some other chlamydial species (Wood, Roshick, & McClarty, 2004), and thus IFN-γ stimulation effectively results in Trp starvation to *C. trachomatis.* The primary regulatory response to Trp starvation in *C. trachomatis* is mediated by a TrpR ortholog, whose Trp-dependent binding to cognate promoter elements represses transcription (Akers & Tan, 2006; Carlson, Wood, Roshick, Caldwell, & McClarty, 2006). This mechanism of regulatory control is presumably limited in *C. trachomatis*, as homologs of genes regulated by TrpR in other bacteria (*e.g. trpF, aroH, aroL*) have not been shown to respond to Trp limitation (Wood et al., 2003).

In many Gram-negative bacteria, such as *Escherichia coli*, *trpR* is monocistronic and distal to the Trp biosynthetic operon. In *C. trachomatis*, TrpR is encoded in an operon, *trpRBA,* which also contains the Trp synthase α- and β- subunits (TrpA and TrpB, respectively), and possesses a 351 base-pair (bp) intergenic region (IGR) that separates *trpR* from *trpBA.* The functional significance of the *trpRBA* IGR is poorly characterized. While a putative TrpR operator sequence was identified in the IGR overlapping an alternative transcriptional origin for *trpBA* (Carlson et al., 2006), TrpR binding was not shown (Akers & Tan, 2006). Based on *in silico* predictions, an attenuator sequence has been annotated within the *trpRBA* IGR (Merino & Yanofsky, 2005), but this has not been thoroughly validated experimentally. Regardless, the IGR is >99% conserved at the nucleotide sequence level across ocular, genital and lymphogranuloma venereum (LGV) serovars of *C. trachomatis*, indicating functional importance (Carlson, Porcella, Mcclarty, & Caldwell, 2005; Seth-Smith et al., 2009; Stephens et al., 1998; Thomson et al., 2008). Therefore, outside of TrpR-mediated repression, the complete detail of *trpRBA* regulation remains poorly elucidated and previous reports have indicated the possibility of more complex mechanisms of regulation (Brinkworth, Wildung, & Carabeo, 2018).

In evaluating alternative regulatory modes of the *trpRBA* operon, an interesting consideration is the pleiotropic effects induced by IFN- γ stimulation of infected cells. IFN- γ is involved in many processes that limit iron and other essential biometals to intracellular pathogens as a component of host nutritional immunity (Cassat & Skaar, 2013; Hood & Skaar, 2012). *Chlamydia* have a strict iron dependence for normal development, evidenced by the onset of persistence following prolonged iron limitation (Raulston, 1997). Importantly, *Chlamydia* presumably acquire iron via vesicular interactions between the chlamydial inclusion and slow-recycling transferrin (Tf)-containing endosomes (Ouellette & Carabeo, 2010). IFN- γ is known to down-regulate transferrin receptor (TfR) expression in both monocytes and epithelial cells with replicative consequences for resident intracellular bacteria (T. F. Byrd & Horwitz, 1993; T. Byrd & Horwitz, 1989; Igietseme, Ananaba, Candal, Lyn, & Black, 1998; Nairz et al., 2008). However, iron homeostasis in *Chlamydia* is poorly understood due to the lack of functionally characterized homologs to iron acquisition machinery that are highly conserved in other bacteria (Pokorzynski, Thompson, & Carabeo, 2017). Only the *ytgABCD* operon, encoding a metal permease, has been clearly linked to iron acquisition (J. D. Miller, Sal, Schell, Whittimore, & Raulston, 2009). Intriguingly, the YtgC (CTL0325) open reading frame (ORF) encodes a N-terminal permease domain fused to a C-terminal DtxR-like repressor domain, annotated YtgR (Akers, HoDac, Lathrop, & Tan, 2011; Thompson, Nicod, Malcolm, Grieshaber, & Carabeo, 2012). YtgR is cleaved from the permease domain during infection and functions as an iron-dependent transcriptional repressor to autoregulate the expression of its own operon (Thompson et al., 2012). YtgR represents the only identified iron-dependent transcriptional regulator in *Chlamydia.* Whether YtgR maintains a more diverse transcriptional regulon beyond the *ytgABCD* operon has not yet been addressed and remains an intriguing question in the context of immune-mediated iron limitation to *C. trachomatis*.

Consistent with the highly reduced capacity of the chlamydial genome, it is likely that *C. trachomatis* has a limited ability to tailor a specific response to each individual stress. In the absence of identifiable homologs for most global stress response regulators in *C. trachomatis*, we hypothesized that primary stress responses to pleiotropic insults may involve mechanisms of regulatory integration, whereby important molecular pathways are co-regulated by stress-responsive transcription factors such that they can be utilized across multiple host-mediated stresses. Here, we report on the unique iron-dependent regulation of the *trpRBA* operon in *Chlamydia trachomatis*. We propose a model of iron-dependent transcriptional regulation of *trpRBA* mediated by the repressor YtgR binding specifically to the IGR, which may have implications for how *C. trachomatis* responds to immunological and environmental insults. Such a mechanism of iron-dependent regulation of Trp biosynthesis has not been previously described in any other prokaryote and adds to the catalog of regulatory models for Trp biosynthetic operons in bacteria. Further, it reveals a highly dynamic mode of regulatory integration within the *trpRBA* operon, employing bipartite control at the transcription initiation and termination steps.

## Results

### Brief iron limitation via 2,2-bipyridyl treatment yields iron-starved, but nonpersistent *Chlamydia trachomatis*

To identify possible instances of regulatory integration between iron and Trp starvation in *C. trachomatis,* we optimized a stress response condition that preceded the development of a characteristically persistent phenotype. We reasoned that in order to effectively identify regulatory integration, we would need to investigate the bacterium under stressed, but not aberrant, growth conditions such that we could distinguish primary stress responses from abnormal growth. To specifically investigate the possible contribution of iron limitation to a broader immunological (*e.g.* IFN-γ-mediated) stress, we utilized the membrane-permeable iron chelator 2,2-bipyridyl (Bpdl), which has the advantage of rapidly and homogeneously starving *C. trachomatis* of iron (Thompson & Carabeo, 2011). We chose to starve *C. trachomatis* serovar L2 of iron starting at 12 hrs post-infection (hpi), or roughly at the beginning of mid-cycle growth. At this point the chlamydial organisms represent a uniform population of replicative RBs that are fully competent, both transcriptionally and translationally, to respond to stress. We treated infected HeLa cell cultures with 100 μM Bpdl or mock for either 6 or 12 hours (hrs) to determine a condition sufficient to limit iron to *C. trachomatis* without inducing hallmark persistent phenotypes. We stained infected cells seeded on glass coverslips with convalescent human sera and analyzed chlamydial inclusion morphology under both Bpdl- and mock-treated conditions by laser point-scanning confocal microscopy (Figure 1A). Following 6 hrs of Bpdl treatment, chlamydial inclusions were largely indistinguishable from mock-treated inclusions, containing a homogeneous population of larger organisms, consistent with RBs in midcycle growth. However, by 12 hrs of Bpdl treatment, the inclusions began to display signs of aberrant growth: they were perceptibly smaller, more comparable in size to 18 hpi, and contained noticeably fewer organisms, perhaps indicating a delay in RB-to-EB differentiation. These observations were consistent with our subsequent analysis of genome replication by quantitative PCR (qPCR; Figure 1B.) At 6 hrs of Bpdl treatment, there was no statistically distinguishable difference in genome copy number when compared to the equivalent mock-treated time-point. However, by 12 hrs of treatment, genome copy number was significantly reduced 4.7-fold in the Bpdl-treated group relative to mock-treatment (*p* = 0.0033). We then assayed the transcript expression of two markers for persistence by reverse transcription quantitative PCR (RT-qPCR): the early gene *euo*, encoding a transcriptional repressor of late-cycle genes (Figure 1C), and the adhesin *omcB*, which is expressed late in the developmental cycle (Figure 1D). Characteristic persistence would display elevated *euo* expression late into infection, and suppressed *omcB* expression throughout development. We observed that at 6 hrs of Bpdl treatment, there was no statistically distinguishable difference in either *euo* or *omcB* expression when compared to the mock-treatment. Still at 12 hrs of Bpdl treatment, *euo* expression was unchanged. However, *omcB* expression was significantly induced following 12 hrs of Bpdl-treatment (*p* = 0.00015). This was unexpected, but we note that *omcB* expression has been shown to vary between chlamydial serovars and species when starved for iron (Pokorzynski et al., 2017). Collectively, these data indicated that 6 hrs of Bpdl treatment was a more suitable timepoint at which to monitor iron-limited stress responses.

**Figure 1.**
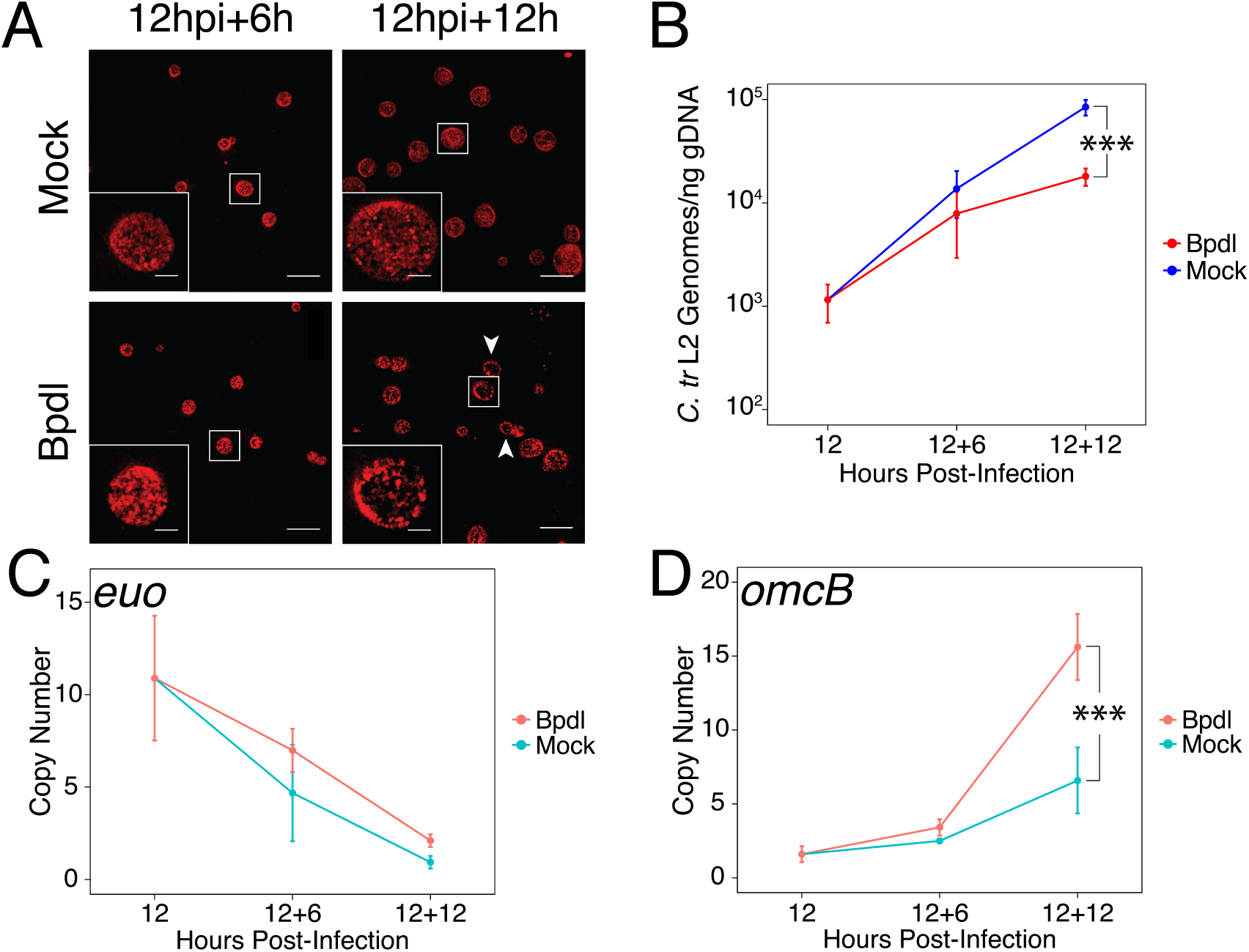
Brief iron limitation via 2,2-bipyridyl treatment precedes the onset of characteristic chlamydial persistence. (*A*) *C. trachomatis* L2-infected HeLa cells were fixed and stained with convalescent human sera to image inclusion morphology by confocal microscopy following Bpdl treatment at the indicated times post-infection. Arrowheads indicate inclusions with visibly fewer organisms in the 12-hour Bpdl-treated condition. Figure shows representative experiment of three biological replicates. Scale bar = 25 μm, Inset scale bar = 5 μm. (*B*) Genomic DNA (gDNA) was harvested from infected HeLa cells at the indicated times post-infection under iron-replete (blue) and - deplete (red) conditions. Chlamydial genome copy number was quantified by qPCR. Chlamydial genome replication is stalled following 12 hours of Bpdl treatment, but not 6. N=2. (*C*) Total RNA was harvested from infected HeLa cells at the indicated times postinfection under iron-replete (teal) and -deplete (orange) conditions. The transcript abundance of hallmark persistence genes *euo* and *(*D*) omcB* were quantified by RT-qPCR and normalized against genome copy number. Only at 12 hours of Bpdl treatment is *omcB* expression significantly affected. N=3 for 12+6, N=2 for 12+12. Statistical significance was determined by One-Way ANOVA followed by post-hoc pairwise t-tests with Bonferroni’s correction for multiple comparisons. * = *p* < 0.05, ** = *p* < 0.01, *** = *p* < 0.005.

We additionally assayed these same metrics following 6 or 12 hrs of Trp starvation by culturing cells in either Trp-replete or Trp-deplete DMEM-F12 media supplemented with fetal bovine serum (FBS) pre-dialyzed to remove amino acids. We observed no discernable change in inclusion morphology out to 12 hrs of Trp starvation (Figure 1 – Figure Supplement 1A), but genome copy numbers were significantly reduced 2.7-fold at this time-point (*p* = 0.00612; Figure 1 – Figure Supplement 1B). The transcript expression of *euo* (Figure 1 – Figure Supplement 1C) and *omcB* (Figure 1 – Figure Supplement 1D) did not significantly change at either treatment duration, but Trp-depletion did result in a 2.0-fold reduction in *omcB* expression, consistent with a more characteristic persistent phenotype. These data therefore also indicated that 6 hrs of treatment would be ideal to monitor non-persistent responses to Trp limitation.

We next sought to determine whether our brief 6-hr Bpdl treatment was sufficient to elicit a transcriptional iron starvation phenotype. We chose to analyze the expression of three previously identified iron-regulated transcripts, *ytgA* (Figure 2A), *ahpC* (Figure 2B) and *devB* (Figure 2C), by RT-qPCR under Bpdl- and mock-treated conditions (Dill, Dessus-Babus, & Raulston, 2009; Thompson & Carabeo, 2011). In addition, we analyzed the expression of one non-iron-regulated transcript, *dnaB* (Figure 2D), as a negative control (Brinkworth et al., 2018). Following 6 hrs of Bpdl treatment, we observed that the transcript expression of the periplasmic iron-binding protein *ytgA* was significantly elevated 1.75-fold relative to the equivalent mock-treated time-point (*p* = 0.0052). However, we did not observe induction of *ytgA* transcript expression relative to the 12 hpi time-point. We distinguish here between *elevated* and *induced* transcript expression, as chlamydial gene expression is highly developmentally regulated. Thus, it can be more informative to monitor longitudinal expression of genes, *i.e.* their induction, as opposed to elevation relative to an equivalent control time-point, which may simply represent a stall in development. While we did not observe induction of *ytgA,* which would be more consistent with an iron-starved phenotype, we reason that this is a consequence of the brief treatment period, and that longer iron starvation would produce a more robust induction of iron-regulated transcripts. Note that the identification of *ytgA* as iron-regulated has only been previously observed following extended periods of iron chelation (J. D. Miller et al., 2009; Raulston et al., 2007; Thompson & Carabeo, 2011). Similarly, we observed that the transcript expression of the thioredoxin *ahpC* was significantly elevated 2.15-fold relative to the equivalent mock-treated time-point (*p* = 0.038) but was not induced relative to the 12 hpi time-point. The transcript expression of *devB*, encoding a 6-phosphogluconolactonase involved in the pentose phosphate pathway, was not observed to significantly respond to our brief iron limitation condition, suggesting that it is not a component of the primary iron starvation stress response in *C. trachomatis*. As expected, the transcript expression of *dnaB*, a replicative DNA helicase, was not altered by our iron starvation condition, consistent with its presumably iron-independent regulation (Brinkworth et al., 2018). Overall, these data confirmed that our 6-hr Bpdl treatment condition was suitable to produce a mild iron starvation phenotype at the transcriptional level, thus facilitating our investigation of iron-dependent regulatory integration.

**Figure 2.**
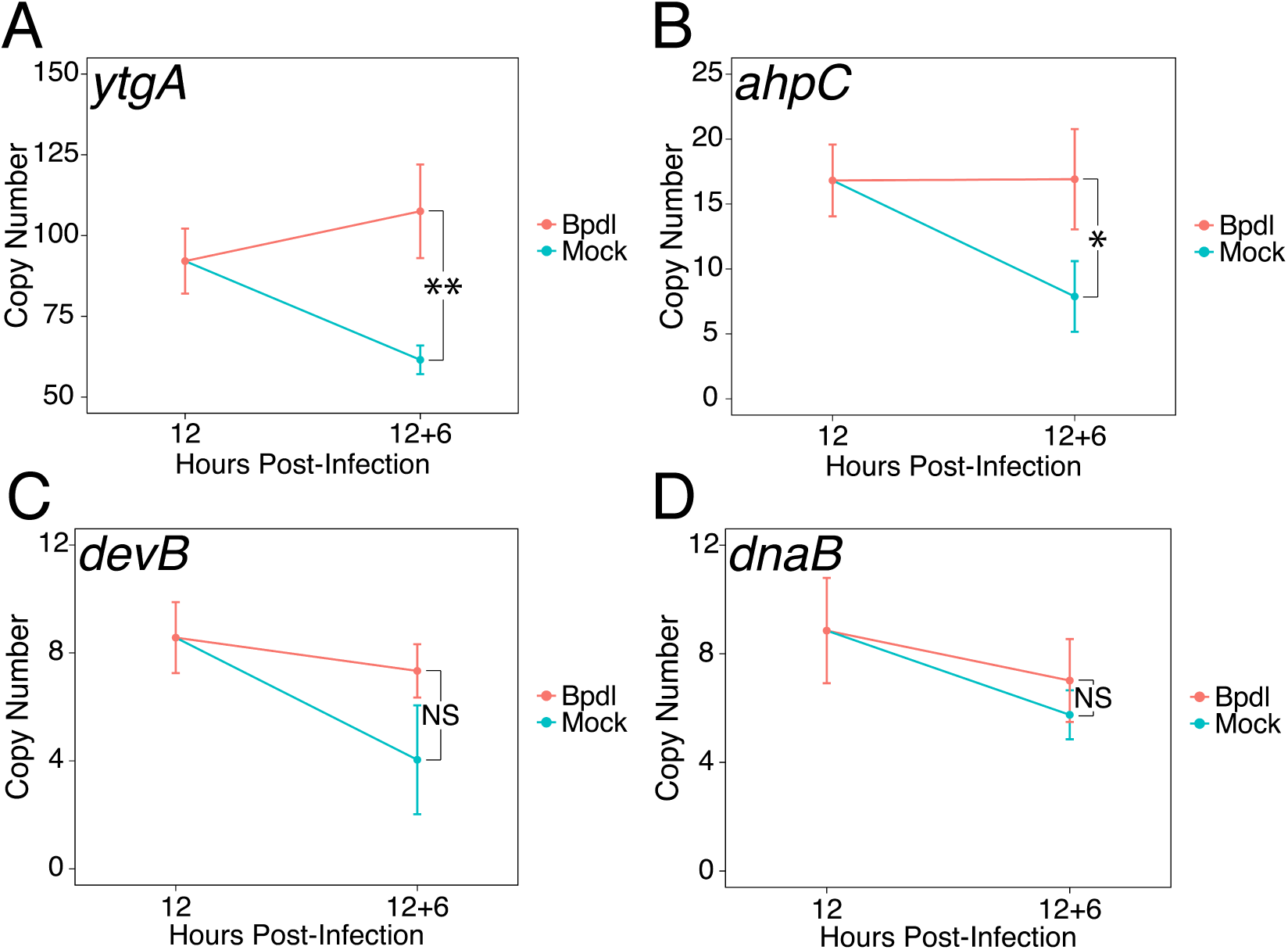
Brief iron limitation condition produces mild iron-starved transcriptional phenotype. (*A*) Total RNA and gDNA was harvested from infected HeLa cells at the indicated times post-infection under iron-replete (teal) and -deplete (orange) conditions. The transcript abundance of iron-regulated *ytgA*, (*B*) *ahpC*, (*C*) *devB* and (*D*) non-iron regulated *dnaB* were quantified by RT-qPCR and normalized against genome copy number. The transcript expression of *ytgA* and *ahpC* were significantly elevated following 6-hour Bpdl treatment, indicative or iron starvation to *C. trachomatis.* N=3. Statistical significance was determined by One-Way ANOVA followed by post-hoc pairwise t-tests with Bonferroni’s correction for multiple comparisons. * = *p* < 0.05, ** = *p* < 0.01, *** = *p* < 0.005.

### Transcript expression of the *trpRBA* operon is differentially regulated by iron in *Chlamydia trachomatis*

Upon identifying an iron limitation condition that produced a relevant transcriptional phenotype while avoiding the onset of persistent development, we aimed to investigate whether the immediate response to iron starvation in *C. trachomatis* would result in the consistent induction of pathways unrelated to iron utilization/acquisition, but nevertheless important for surviving immunological stress. The truncated Trp biosynthetic operon, *trpRBA* (Figure 3A), has been repeatedly linked to the ability of genital and LGV serovars (D-K and L1-3, respectively) of *C. trachomatis* to counter IFN-γ-mediated stress. This is due to the capacity of the chlamydial Trp synthase in these serovars to catalyze the β synthase reaction, *i.e.* the condensation of indole to the amino acid serine to form Trp (Fehlner-Gardiner et al., 2002). In the presence of exogenous indole, *C. trachomatis* is therefore able to biosynthesize Trp such that it can prevent the development of IFN-γ-mediated persistence. Correspondingly, the expression of *trpRBA* is highly induced following IFN-γ stimulation of infected cells (Belland et al., 2003; Østergaard et al., 2016). These data have historically implicated Trp starvation as the primary mechanism by which persistence develops in *C. trachomatis* following exposure to IFN-γ. However, these studies have routinely depended on prolonged treatment conditions that monitor the terminal effect of persistent development, as opposed to the immediate molecular events which may have important roles in the developmental fate of *Chlamydia*. As such, these studies may have missed the contribution of other IFN-γ-stimulated insults such as iron limitation.

**Figure 3.**
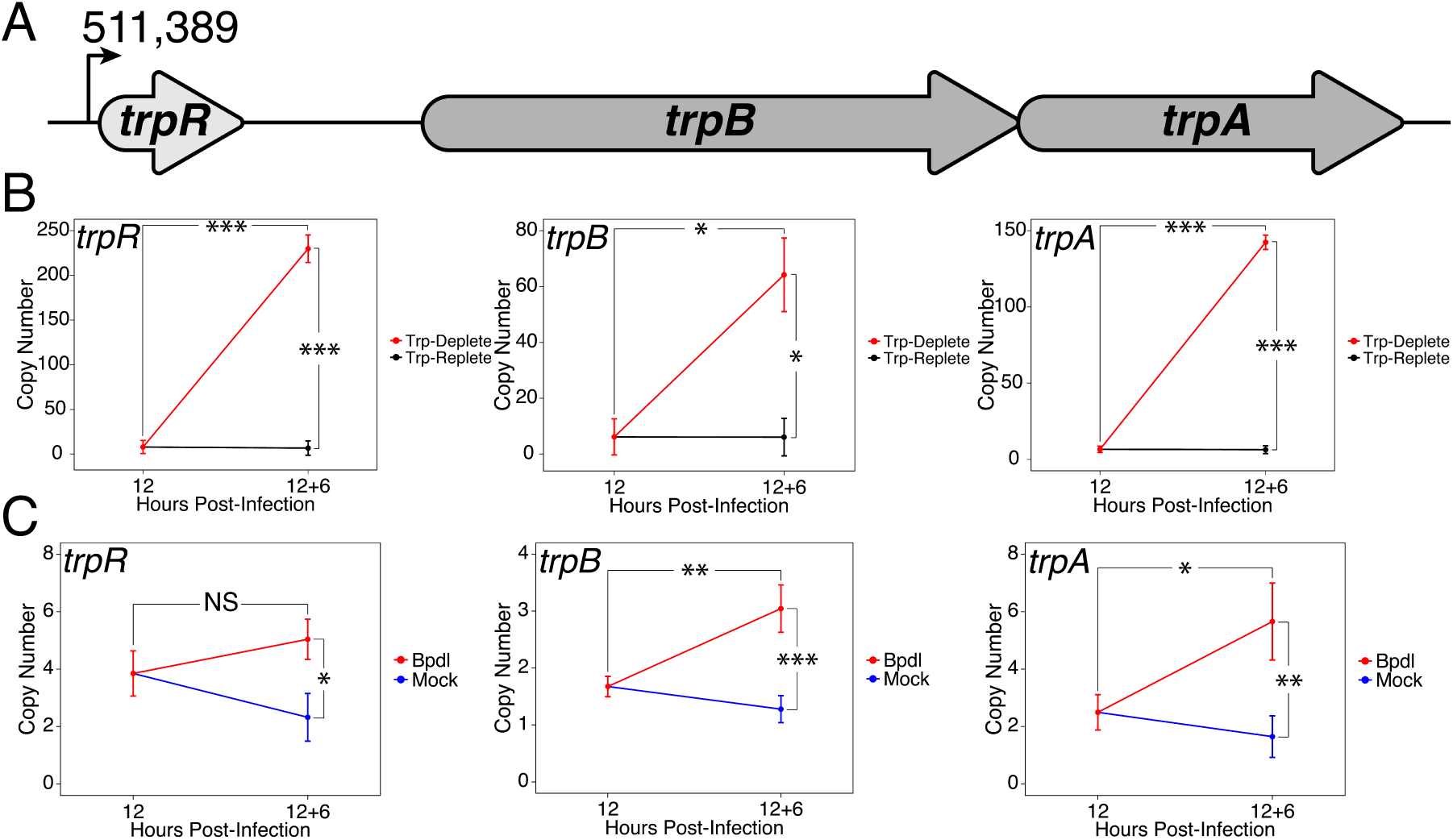
Expression of the *trpRBA* operon in *C. trachomatis* is differentially regulated by brief iron limitation. (*A*) Cartoon depiction of the *trpRBA* operon (drawn to scale) with the primary transcriptional start site upstream of *trpR* annotated. (*B*) Total RNA and gDNA were harvested from infected HeLa cells at the indicated times post-infection under Trp-replete (black) and -deplete (red) conditions. The transcript expression of *trpRBA* operon was quantified by RT-qPCR and normalized against genome copy number. All three ORFs are significantly induced relative to 12 hpi following Trp starvation. N=2. (*C*) Total RNA and gDNA were harvested from infected HeLa cells at the indicated times post-infection under iron-replete (blue) and -deplete (red) conditions. The transcript expression of *trpRBA* operon was quantified by RT-qPCR and normalized against genome copy number. Only *trpB* and *trpA* expression was significantly induced relative to 12 hpi. N=3. Statistical significance was determined by One-Way ANOVA followed by post-hoc pairwise t-tests with Bonferroni’s correction for multiple comparisons. * = *p* < 0.05, ** = *p* < 0.01, *** = *p* < 0.005.

To decouple Trp limitation from iron limitation and assess their relative contribution to regulating a critical pathway for responding to IFN-γ-mediated stress, we monitored the transcript expression of the *trpRBA* operon under brief Trp or iron starvation by RT-qPCR. When starved for Trp for 6 hrs, we observed that the expression of *trpR*, *trpB* and *trpA* were all significantly induced greater than 10.5-fold relative to 12 hpi (*p* = 0.00077, 0.025 and 9.7e-5, respectively; Figure 3B). All three ORFs were also significantly elevated relative to the equivalent mock-treated time-point (*p* = 0.00076, 0.025 and 9.7e-5, respectively). This result was surprising with respect to the relative immediacy of operon induction in response to our Trp starvation protocol, confirming the relevant Trp-starved transcriptional phenotype. To induce Trp-deprived persistence in *C. trachomatis*, many laboratories rely on compounded techniques of IFN-γ pre-treatment to deplete host Trp pools in conjunction with culturing in Trp-depleted media, among other strategies. While the phenotypic end-point differs here, it is nonetheless interesting to note that only 6 hrs of media replacement is sufficient to markedly up-regulate *trpRBA* expression. This suggests that *C. trachomatis* has a highly attuned sensitivity to even moderate changes in Trp levels.

We next performed the same RT-qPCR analysis on the expression of the *trpRBA* operon in response to 6 hrs of iron limitation via Bpdl treatment (Figure 3C). While we observed that the transcript expression of all three ORFs was significantly elevated at least 2.1-fold relative to the equivalent mock-treated time-point (*p* = 0.015, 0.00098 and 0.0062, respectively), we made the intriguing observation that only the expression of *trpB* and *trpA* was significantly induced relative to 12 hpi (*p* = 0.00383 and 0.0195, respectively). The significant induction of *trpBA* expression, but not *trpR* expression, suggested that *trpBA* are specifically regulated by iron availability. This result is consistent with a recent survey of the iron-regulated transcriptome in *C. trachomatis* by RNA-Sequencing, which also reported that iron-starved *Chlamydia* specifically up-regulate *trpBA* expression in the absence of altered *trpR* expression (Brinkworth et al., 2018). Our results expand on this finding by providing a more detailed investigation into the specific profile of this differential regulation of *trpRBA* in response to iron deprivation. Taken together, these findings demonstrate that an important stress response pathway, the *trpRBA* operon, is regulated by the availability of both Trp and iron, consistent with the notion that the pathway may be cooperatively regulated to respond to various stress conditions. Notably, iron-dependent regulation of Trp biosynthesis has not been previously documented in other prokaryotes.

### Specific iron-regulated expression of *trpBA* originates from a novel alternative transcriptional start site within the *trpRBA* intergenic region

We hypothesized that the specific iron-related induction of *trpBA* expression relative to *trpR* expression may be attributable to an iron-regulated alternative transcriptional start site (alt. T<u>S</u>S) downstream of the *trpR* ORF. Indeed, a previous study reported the presence of an alt. TSS in the *trpRBA* IGR, located 214 nucleotides upstream of the *trpB* translation start position (Carlson et al., 2006). However, a parallel study could not identify a TrpR binding site in the *trpRBA* IGR (Akers & Tan, 2006). We reasoned that a similar alt. TSS may exist in the IGR that controlled the iron-dependent expression of *trpBA.* We therefore performed <u>R</u>apid <u>A</u>mplification of <u>5</u>’-<u>c</u>DNA <u>E</u>nds (5’-RACE) on RNA isolated from *C. trachomatis* L2-infected HeLa cells using the SMARTer 5’/3’ RACE Kit workflow (Takara Bio). Given the low expression of the *trpRBA* operon during normal development, we utilized two sequential gene-specific amplification steps (nested 5’-RACE) to identify 5’ cDNA ends in the *trpRBA* operon. These nested RACE conditions resulted in amplification that was specific to infected-cells (Figure 4 – Figure Supplement 1A). Using this approach, we analyzed four conditions: 12 hpi, 18 hpi, 12 hpi + 6 hrs of Bpdl treatment, and 12 hpi + 6 hrs of Trp-depletion (Figure 4A). We observed three RACE products that migrated with an apparent size of 1.5, 1.1 and 1.0 kilobases (kb). At 12 and 18 hpi, all three RACE products exhibited low abundance, even following the nested PCR amplification. This observation was consistent with the expectation that the expression of the *trpRBA* operon is very low under normal, iron and Trp-replete conditions. However, we note that the 6-hr difference in development did appear to alter the representation of the 5’ cDNA ends, which may suggest a stage-specific promoter utilization within the *trpRBA* operon. In our Trp starvation condition, we observed an apparent increase in the abundance of the 1.5 kb RACE product, which was therefore presumed to represent the primary TSS upstream of *trpR,* at nucleotide position 511,389 (*C. trachomatis* L2 434/Bu). Interestingly, the 1.0 kb product displayed a very similar apparent enrichment following Bpdl treatment, suggesting that this RACE product represented a specifically iron-regulated TSS. Both the 1.5 and 1.0 kb RACE products were detectable in the Trp-depleted and iron-depleted conditions, respectively, during the primary RACE amplification, consistent with their induction under these conditions (Figure 4 – Figure Supplement 1B).

**Figure 4.**
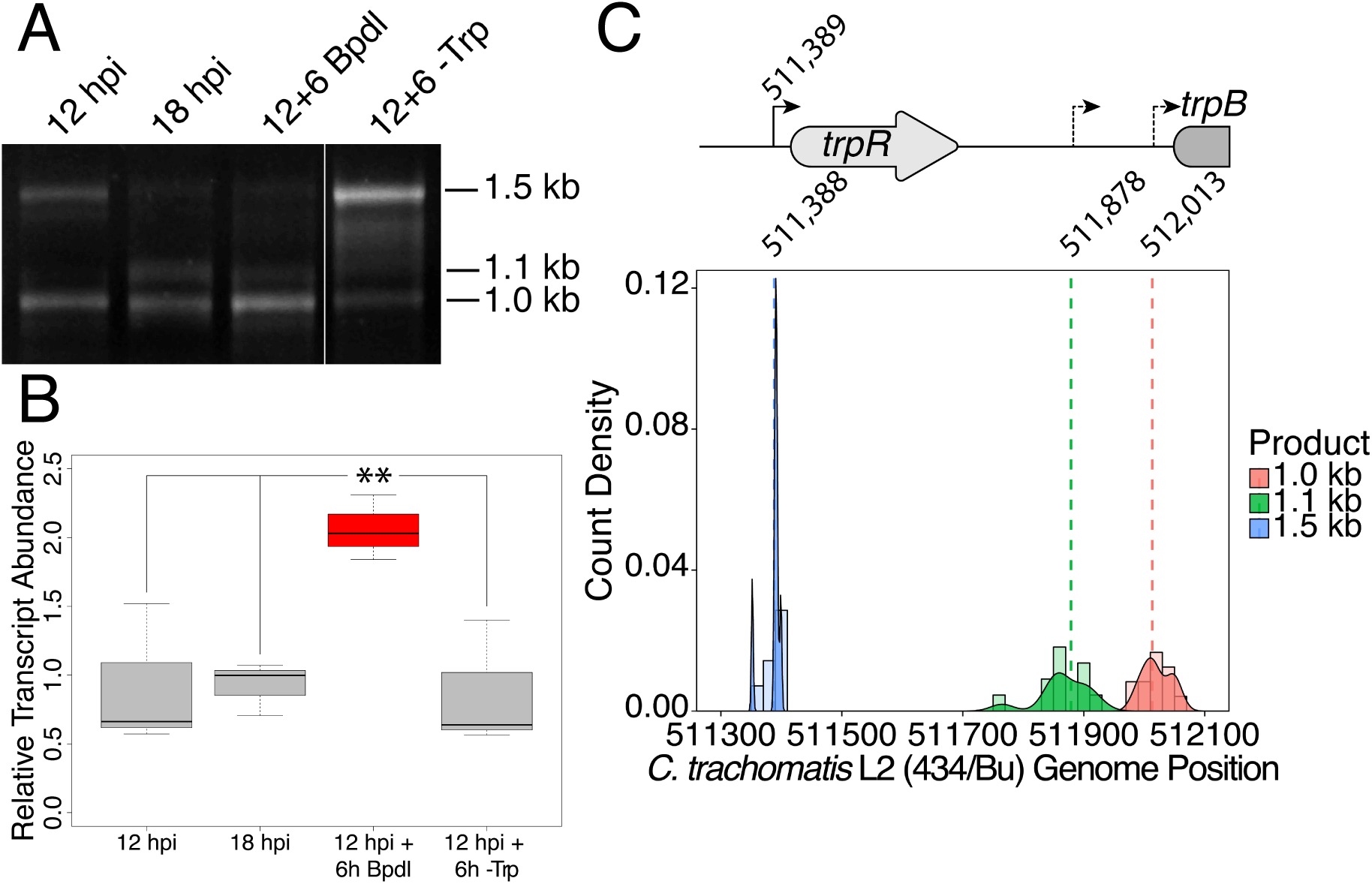
Iron-dependent induction of *trpBA* expression initiates within the *trpRBA* intergenic region from a novel alternative transcriptional start site. (*A*) Total RNA was harvested from infected HeLa cells at the indicated times post-infection to examine iron-dependent and Trp-dependent changes in the 5’-cDNA profile of the *trpRBA* operon by Rapid Amplification of 5’ cDNA Ends (5’-
RACE). RACE products were separated on an agarose gel, revealing three distinct and specific bands with apparent sizes of 1.5, 1.1 and 1.0 kb. Trp depletion led to the apparent enrichment of the 1.5 kb product, while Bpdl treatment produced a similarly enriched 1.0 kb RACE product. Figure shows representative experiment of three biological replicates. (*B*) To confirm that iron-dependent induction of *trpBA* could originate from alternative transcription initiation, RT-qPCR was performed on 5’-RACE total RNA to quantify the abundance of *trpB* transcripts relative to *trpR.* Only under iron-limited conditions were *trpB* transcripts enriched relative to *trpR.* N=3. Statistical significance determined by One-way ANOVA followed by post-hoc pairwise t-tests. * = *p* < 0.05, ** = *p* < 0.01, *** = *p* < 0.005. (*C*) The nucleotide position of the 5’ cDNA ends generated from RACE were mapped to the *C. trachomatis* L2 434/Bu genome by nucleotide BLAST. Figure displays histogram (semi-continuous; bin width=20) and overlaid density plot (continuous) distribution of 5’ nucleotide positions generated from each 5’-RACE product. The dotted line represents the weighted mean of the distribution, as indicated by the integer value above each line. The identified alt. TSSs are depicted on the *trpRBA* operon (drawn to scale) above the plot. N=3.

If iron depletion was inducing *trpBA* expression independent of *trpR,* we reasoned that we would observe specific enrichment of *trpB* sequences in our 5’-RACE cDNA samples relative to *trpR* sequences. We again utilized RT-qPCR to quantify the abundance of *trpB* transcripts relative to *trpR* transcripts in the 5’-RACE total RNA samples (Figure 4B). In agreement with our model, only under iron starved conditions did we observe a significant enrichment of *trpB* relative to *trpR* (*p* < 0.01). Additionally, we observed that at 12 and 18 hpi in iron-replete conditions, the ratio of *trpB* to *trpR* was approximately 1.0, suggesting non-preferential basal expression across the three putative TSSs. Another factor contributing to this ratio is the synthesis of the full-length *trpRBA* polycistron. In support of this, the *trpB* to *trpR* ratio remained near 1.0 under the Trp-starved condition, which would be expected during transcription read-through of the whole operon. The apparent lack of preferential promoter utilization as described above could be attributed to the relatively low basal expression of the operon at 12 and 18 hpi under Trp- and iron-replete conditions, thus precluding quantitative detection of differential promoter utilization in this assay.

To determine the specific location of the 5’ cDNA ends within the *trpRBA* operon, we isolated the 5’-RACE products across all conditions by gel extraction and cloned the products into the pRACE vector supplied by the manufacturer. We then sequenced the ligated inserts and BLASTed the sequences against the *C. trachomatis* L2 434/Bu genome to identify the location of the 5’-most nucleotides (Figure 4C). These data are displayed as a statistical approximation of the genomic regions most likely to be represented by the respective 5’-RACE products in both histogram (semi-continuous) and density plot (continuous) format (See Supplementary File 1 for a description of all mapped 5’-RACE products). As expected, the 1.5 kb product mapped in a distinct and tightly grouped peak near the previously annotated *trpR* TSS, with the mean and modal nucleotide being 511,388 and 511,389, respectively (Figure 4 – Figure Supplement 2A). Surprisingly, we found that neither the 1.1 or 1.0 kb RACE product mapped to the previously reported alt. TSS in the *trpRBA* IGR, at position 511,826. Instead, we observed that the 1.1 kb product mapped on average to nucleotide position 511,878, with the modal nucleotide being found at 511,898 (Figure 4 – Figure Supplement 2B). The 1.0 kb product mapped with a mean nucleotide position of 512,013, with the modal nucleotide being 512,005 (Figure 4 – Figure Supplement 2C), only 35 bases upstream of the *trpB* coding sequence. Interestingly, the 1.0 kb product mapped to a region of the *trpRBA* IGR flanked by consensus σ^66^ -10 and -35 promoter elements, found at positions 512,020-5 and 511,992-7, respectively (Ricci, Ratti, & Scarlato, 1995). These data collectively pointed toward the 1.0 kb 5’-RACE product representing a novel, iron-regulated alt. TSS and *bona fide* σ^66^-dependent promoter element that allows for the specific iron-dependent expression of *trpBA.*

### YtgR specifically binds to the *trpRBA* intergenic region in an operator-dependent manner to repress transcription of *trpBA*

As the only known iron-dependent transcriptional regulator in *Chlamydia*, we hypothesized that YtgR may regulate the iron-dependent expression of *trpBA* from the putative promoter element we characterized by 5’-RACE. Using bioinformatic sequence analysis, we investigated whether the *trpRBA* IGR contained a candidate YtgR operator sequence. By local sequence alignment of the putative YtgR operator sequence (Akers et al., 2011) and the *trpRBA* IGR, we identified a high-identity alignment (76.9% identity) covering 67% of the putative operator sequence (Figure 5A). Interestingly, this alignment mapped to the previously identified palindrome suspected to have operator functionality (Carlson et al., 2006). By global sequence alignment of the YtgR operator to the palindromic sequence, an alignment identical to the local alignment was observed, which still displayed relatively high sequence identity (43.5% identity). We hypothesized that this sequence functioned as an YtgR operator, despite being located 184 bp upstream of the *trpBA* alt. TSS.

**Figure 5.**
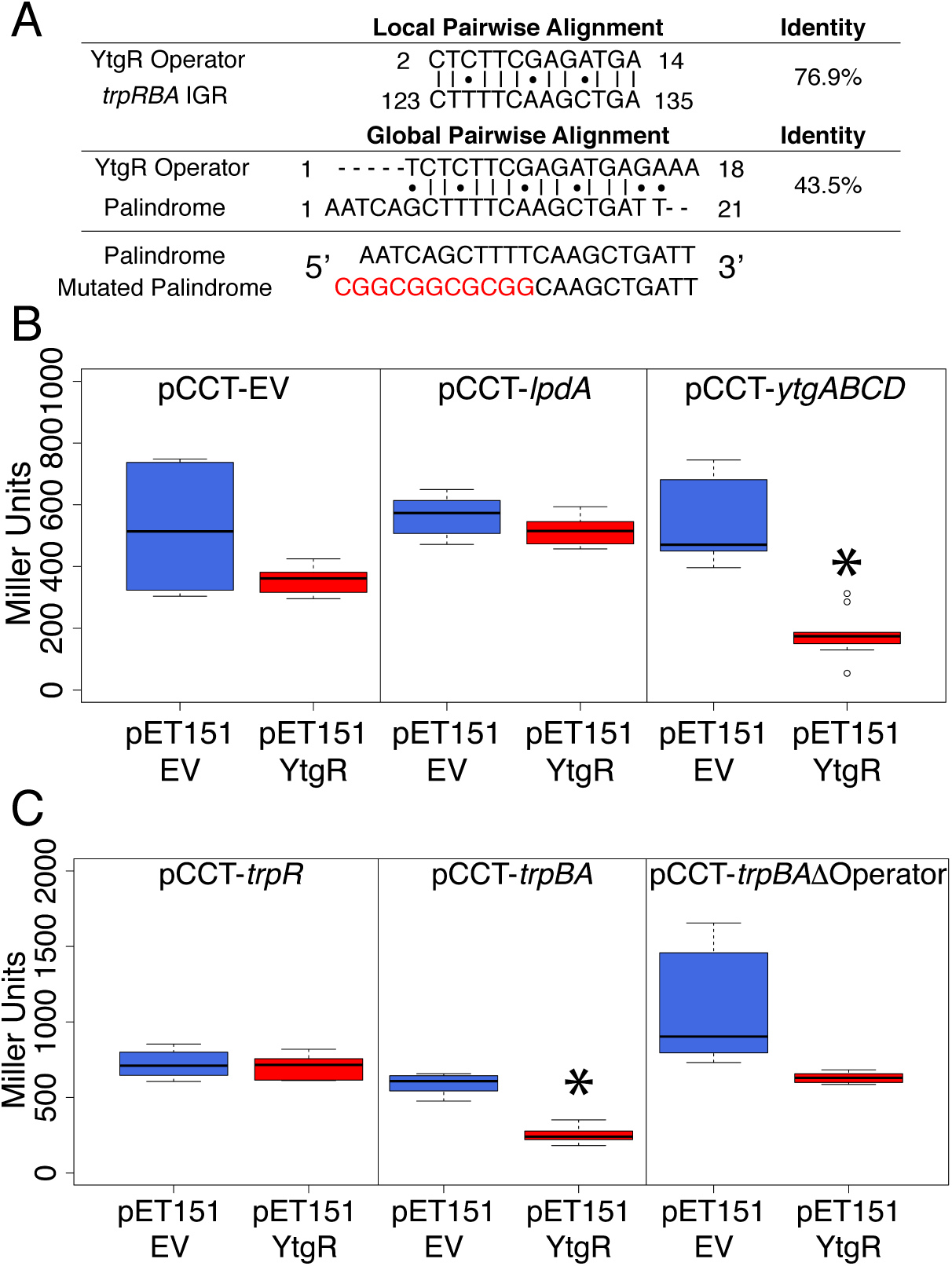
Ectopically expressed YtgR domain is capable of binding the putative *trpBA* promoter element in an operator-specific manner to repress transcription in a heterologous system. (*A*) Identification of putative YtgR operator sequence by local and global nucleotide sequence alignment using EMBOSS Water and Needle algorithms, respectively, to align the previously identified YtgR operator to both the *trpRBA* IGR and palindromic candidate sequence. The palindrome was then mutated in our YtgR repression assay as depicted to abolish palindromicity and AT-richness. (*B*) Ectopic expression of YtgR significantly represses β-galactosidase activity only from the promoter of its own operon, *ytgABCD,* and not from an empty vector or another iron-regulated but presumably non-YtgR targeted promoter, *lpdA.* N = 2 or 3. (*C*) Expression of recombinant YtgR represses β-galactosidase activity from the putative *trpBA* promoter element, but not the *trpR* promoter, and this repression is dependent on the unaltered operator sequence identified in Fig. 5A. N = 2 or 3. Statistical significance determined by two-sided unpaired Student’s *t*-test with Welch’s correction for unequal variance. * = *p* < 0.05, ** = *p* < 0.01, *** = *p* < 0.005.

To investigate the ability of YtgR to bind and repress transcription from the putative *trpBA* promoter, we implemented a heterologous two-plasmid assay that reports on YtgR repressor activity as a function of β-galactosidase expression (Thompson et al., 2012). In brief, a candidate DNA promoter element was cloned into the pCCT expression vector between an arabinose-inducible pBAD promoter and the reporter gene *lacZ*. This plasmid was co-transformed into BL21 (DE3) *E. coli* along with an IPTG-inducible pET151 expression vector with (pET151-YtgR) or without (pET151-EV) the C-terminal 139 amino acid residues of CTL0325 (YtgC). Note that we have previously demonstrated that this region is a functional iron-dependent repressor domain (Thompson et al., 2012). To verify the functionality of this assay, we determined whether ectopic YtgR expression could repress pCCT reporter gene expression in the presence of three candidate DNA elements: a no-insert empty vector (pCCT-EV), the putative promoter element for *C. trachomatis IpdA* (pCCT-*lpdA*), and the promoter region of the *ytgABCD* operon (pCCT-*ytgABCD*; Figure 5B). As expected, from the pCCT-EV reporter construct, ectopic YtgR expression did not significantly reduce the activity of β-galactosidase. Additionally, reporter gene expression from pCCT-*lpdA*, containing the promoter of iron-regulated *lpdA* (Brinkworth et al., 2018), which is not known to be YtgR-regulated, was not affected by ectopic expression of YtgR. This demonstrated that the assay can discriminate between the promoter elements of iron-regulated genes and *bona fide* YtgR targeted promoters. Indeed, in the presence of pCCT-*ytgABCD*, induction of YtgR expression produced a significant decrease in β-galactosidase activity (*p* = 0.03868) consistent with its previously reported auto-regulation of this promoter (Thompson et al., 2012).

Using this same assay, we then inserted into the pCCT reporter plasmid 1) the *trpR* promoter element (pCCT-*trpR*), 2) the putative *trpBA* promoter element represented by the IGR (pCCT-*trpBA*), and 3) the same putative *trpBA* promoter element with a mutated YtgR operator sequence that was diminished for both palindromicity and A-T richness, two typical features of prokaryotic promoter elements (pCCT-*trpBA*ΔOperator; Figure 5C) (Schmitt, 2002; Tao, Boydt, & Murphy, 1992). When YtgR was ectopically expressed in the pCCT-*trpR* background, we observed no statistically distinguishable change in β-galactosidase activity. However, in the pCCT-*trpBA* background, ectopic YtgR expression significantly reduced β-galactosidase activity at levels similar to those observed in the pCCT-*ytgABCD* background (*p* = 0.01219). This suggested that YtgR was capable of binding to the *trpBA* promoter element specifically. Interestingly, this repression phenotype was abrogated in the pCCT-*trpBA*ΔOperator background, where we observed no statistically meaningful difference in β-galactosidase activity. We subsequently addressed whether the region of the *trpRBA* IGR containing the YtgR operator site was sufficient to confer YtgR repression in this assay (Figure 5 – Figure Supplement 1). We cloned three fragments of the *trpRBA* IGR into the pCCT reporter plasmid: the first fragment represented the 5’- end of the IGR containing the operator site at the 3’-end (pCCT-IGR1), the second fragment represented a central region of the IGR containing the operator site at the 5’- end (pCCT-IGR2), and the third fragment represented the 3’-end of the IGR and did not contain the operator site (pCCT-IGR3). Surprisingly, we observed that none of these fragments alone were capable of producing a significant repression phenotype in our reporter system. This finding indicated that while the operator site was necessary for YtgR repression, it alone was not sufficient. Together, these data indicated that YtgR could bind to the *trpBA* promoter element and that this binding was dependent upon an intact AT-rich palindromic sequence, likely representing an YtgR operator, but that further structural elements in the *trpRBA* IGR may be necessary for repression. Nonetheless, we demonstrated the existence of a functional YtgR binding site that conferred iron-dependent transcriptional regulation to *trpBA,* independent of the major *trpR* promoter.

### Transcripts initiated at the primary *trpR* promoter terminate at the YtgR operator site

We hypothesized that YtgR binding at the *trpRBA* YtgR operator site may disadvantage the processivity of RNAP reading-through the IGR from the upstream *trpR* promoter. Similar systems of RNAP read-through blockage have been reported; the transcription factor Reb1p “roadblocks” RNAPII transcription read-through in yeast by promoting RNAP pausing and subsequent degradation (Colin et al., 2014). To investigate this question, we first returned to RNA-Sequencing data we generated to define the immediate iron-dependent transcriptional regulon in *C. trachomatis* (Brinkworth et al., 2018). Using data obtained from *C. trachomatis*-infected HeLa cells at 12 hpi + 6h mock or Bpdl treatment, we mapped the sequenced reads in batch across three biological replicates to the *C. trachomatis* L2 434/Bu genome (NC_010287) which we modified to include annotations for non-operonic IGRs. This coverage map indicated that under Bpdl-treated conditions, a higher proportion of reads mapped to the *trpRBA* IGR (IGR_trpB) relative to mock-treatment (Figure 6A). However, a notable increase in reads mapping to the upstream *trpR* CDS was not observed, suggesting that under iron replete conditions, transcripts originating from the primary *trpR* promoter may be terminated before reading through the IGR. Note that this is consistent with the original observation that *trpR* is not differentially expressed under iron-limited conditions in this RNA-Seq dataset (Brinkworth et al., 2018). We additionally investigated the abundance of reads mapping to the IGRs upstream of *euo* (IGR_euo; not iron-regulated, not YtgR-regulated) and *lpdA* (IGR_lpdA; iron-regulated, not YtgR-regulated), and were unable to observe a similar increase in read coverage at these IGRs following Bpdl treatment, indicating that the increased coverage at the *trpRBA* IGR was specific (Figure 6 – Figure Supplement 1A-B). The absolute number of reads mapping to the *trpRBA* IGR under these conditions was very low relative to either upstream or downstream CDS, implying that the terminated transcript species are rare. We therefore turned to more sensitive and quantitative methods to interrogate possible transcript termination within the *trpRBA* IGR.

**Figure 6.**
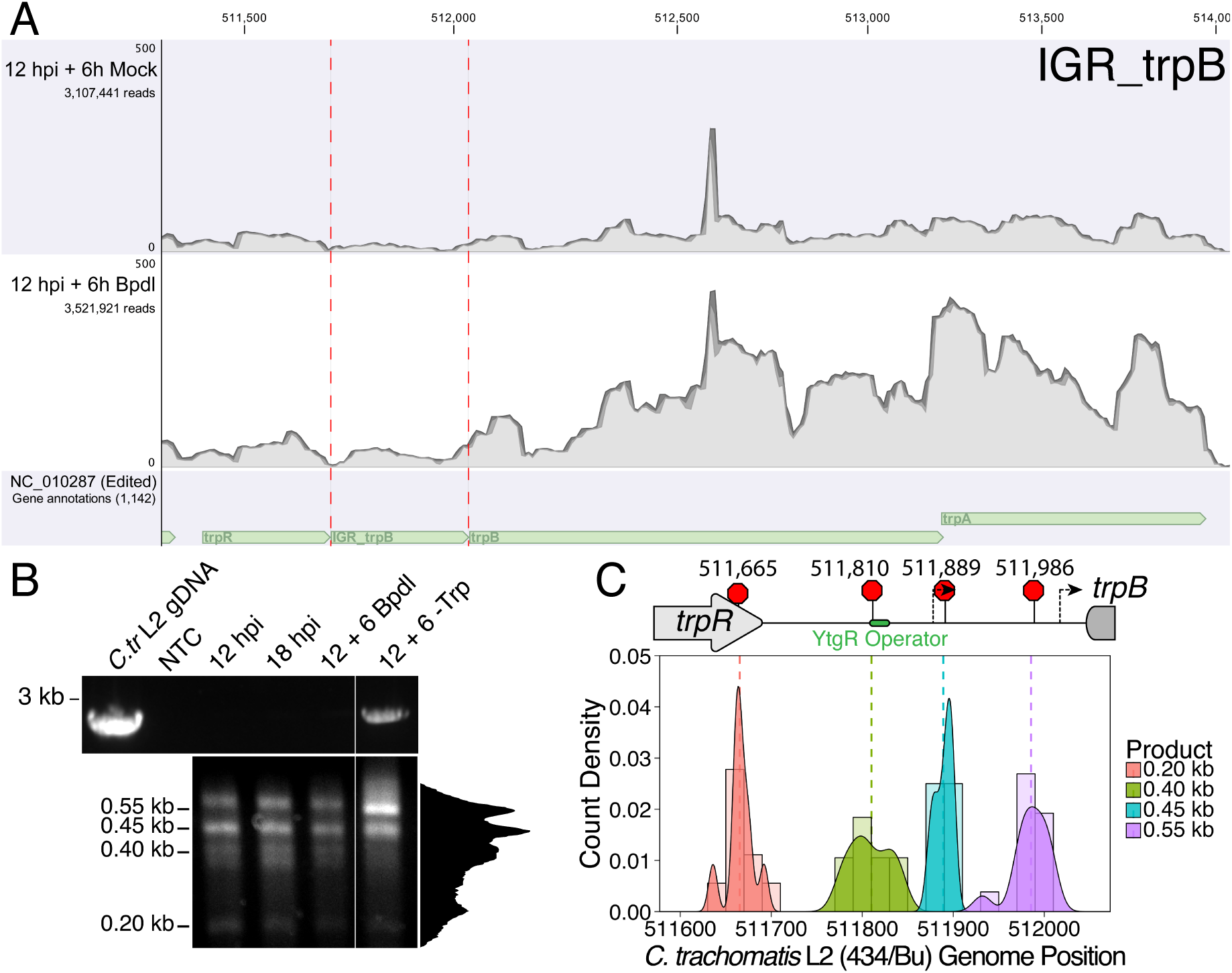
Transcription from the primary *trpR* promoter terminates in the *trpRBA* IGR region, notably at the YtgR operator site. (*A*) Coverage map of RNA-Sequencing reads mapped to the *C. trachomatis* L2/434 Bu genome (NC_010287) edited to contain annotations for non-operonic IGRs. Read coverage at the *trpRBA* IGR (IGR_trpB) is increased following Bpdl treatment, but *trpR* read coverage is not similarly increased. (*B*) Total RNA was harvested from *C. trachomatis*-infected HeLa cells to analyze 3’- cDNA landscape downstream of the *trpR* promoter. The top panel displays representative RT-PCR of full-length *trpRBA* message across experimental conditions (NTC = No Template Control). Bottom panel depicts electrophoresed 3’-RACE products and estimated sizes. Intensity plot to the right of image was generated using the Fiji Dynamic ROI Profiler plugin to monitor intensity across the 18 hpi condition. Note the presence of four distinct peaks, corresponding to each 3’-RACE product. N=3. (*C*) 3’- RACE products were sequenced and mapped to *C. trachomatis* L2 434/Bu genome by nucleotide BLAST. Figure displays histogram (semi-continuous; bin width=20) and overlaid density plot (continuous) distribution of 3’ nucleotide positions generated from each 3’-RACE product. The dotted line represents the weighted mean of the distribution, as indicated by the integer value above each line. The identified alt. TTSs are depicted on the *trpRBA* operon (drawn to scale) above the plot. The 0.40 kb RACE product mapped to a region overlapping the predicted YtgR operator site. N=.

To identify transcription termination sites (T<u>T</u>Ss) in the *trpRBA* operon in *C. trachomatis,* we utilized 3’-RACE to map the 3’-ends of transcripts using gene-specific primers within the *trpR* CDS (Figure 6B; bottom). We again utilized two RACE amplification cycles to generate distinct, specific bands suitable for isolation and sequencing (Figure 6 – Figure Supplement 2B-C). By gel electrophoresis of the 3’- RACE products, we observed the appearance of four distinct bands that migrated with an apparent size of 0.55, 0.45, 0.40 and 0.20 kb. In our Trp-depleted condition, we observed only a very weak amplification of the 2.5 – 3 kb full-length *trpRBA* message by 3’-RACE (Figure 6 – Figure Supplement 2A). However, we did observe it across all replicates. To confirm that the full-length product was specific to the Trp-deplete treatment, we amplified the *trpRBA* operon by RT-PCR from the 3’-RACE total RNA (Figure 6B; top). As expected, only in the Trp-deplete sample did we observe robust amplification of the full-length *trpRBA* message. We note however that image contrast adjustment reveals a very weak band present in all experimental samples. In accordance with the RNA-Sequencing data, 3’-RACE demonstrated the presence of unique transcription termination events in the *trpRBA* operon.

To identify the specific TTS locations, we gel extracted the four distinct 3’-RACE bands across all conditions and cloned them into the pRACE sequencing vector as was done for the 5’-RACE experiments. We then sequenced the inserted RACE products and mapped them to the *C. trachomatis* L2 434/Bu genome (Figure 6B). This revealed a highly dynamic TTS landscape within the *trpRBA* IGR, which has not previously been investigated (For a full description of mapped 3’-RACE products, see Supplementary File 2). The 0.20 kb RACE product mapped to the 3’-end of the *trpR* CDS, with a mean nucleotide position of 511,665 and a modal nucleotide position of 511,667 (Figure 6 – Figure Supplement 3A). Contrastingly, the other three 3’-RACE products did not map in such a way so as to produce specific, unambiguous modal peaks. Instead, their distribution was broader and more even, with only a few nucleotide positions mapping more than once. Accordingly, the 0.45 kb product mapped with an average nucleotide position of 511,889, just downstream of the 1.1 kb 5’-RACE product (Figure 6 – Figure Supplement 3C), while the 0.55 kb product mapped with an average nucleotide position of 511,986, upstream of the 1.0 kb 5’-RACE product (Figure 6 – Figure Supplement 3D). Interestingly, the 0.40 kb product mapped to a region directly overlapping the putative YtgR operator site, with a mean nucleotide position of 511,811 (Figure 6 – Figure Supplement 3B). We therefore reasoned that this putative TTS may have an iron-dependent function.

### Iron limitation promotes transcription read-through at the YtgR operator site

Given the observation that transcripts terminated at the YtgR operator site, we hypothesized that YtgR binding may promote transcription termination at this locus. Conversely, we hypothesized that inactivating YtgR DNA-binding by Bpdl treatment would allow transcription to read through the YtgR operator site. To quantitatively analyze the possibility that iron-depletion, and thus dissociation of YtgR, may facilitate transcription read-through at the operator site, we utilized RT-qPCR to monitor the abundance of various amplicons across the *trpRBA* operon (Figure 7A). We quantified these data in relation to a “read-through” normalization amplicon that, based on 5’- and 3’-RACE data, should only be represented when the full-length *trpRBA* message is transcribed (Figure 7A). The representation of a specific mRNA species relative to the full-length transcript should therefore be interpretable through a simple ratio of the experimental amplicon to the “read-through” amplicon. If an mRNA species is poorly represented relative to the full-length transcript, the ratio should be approximately 1.0. Conversely, if that species is over-represented relative to the full-length transcript, the ratio should exceed 1.0 (Figure 7B; left). Therefore, as each amplicon is increasingly represented as a part of the full-length transcript, the ratio of the specific amplicon to the normalization amplicon should approach 1.0 (Figure 7B; right).

**Figure 7.**
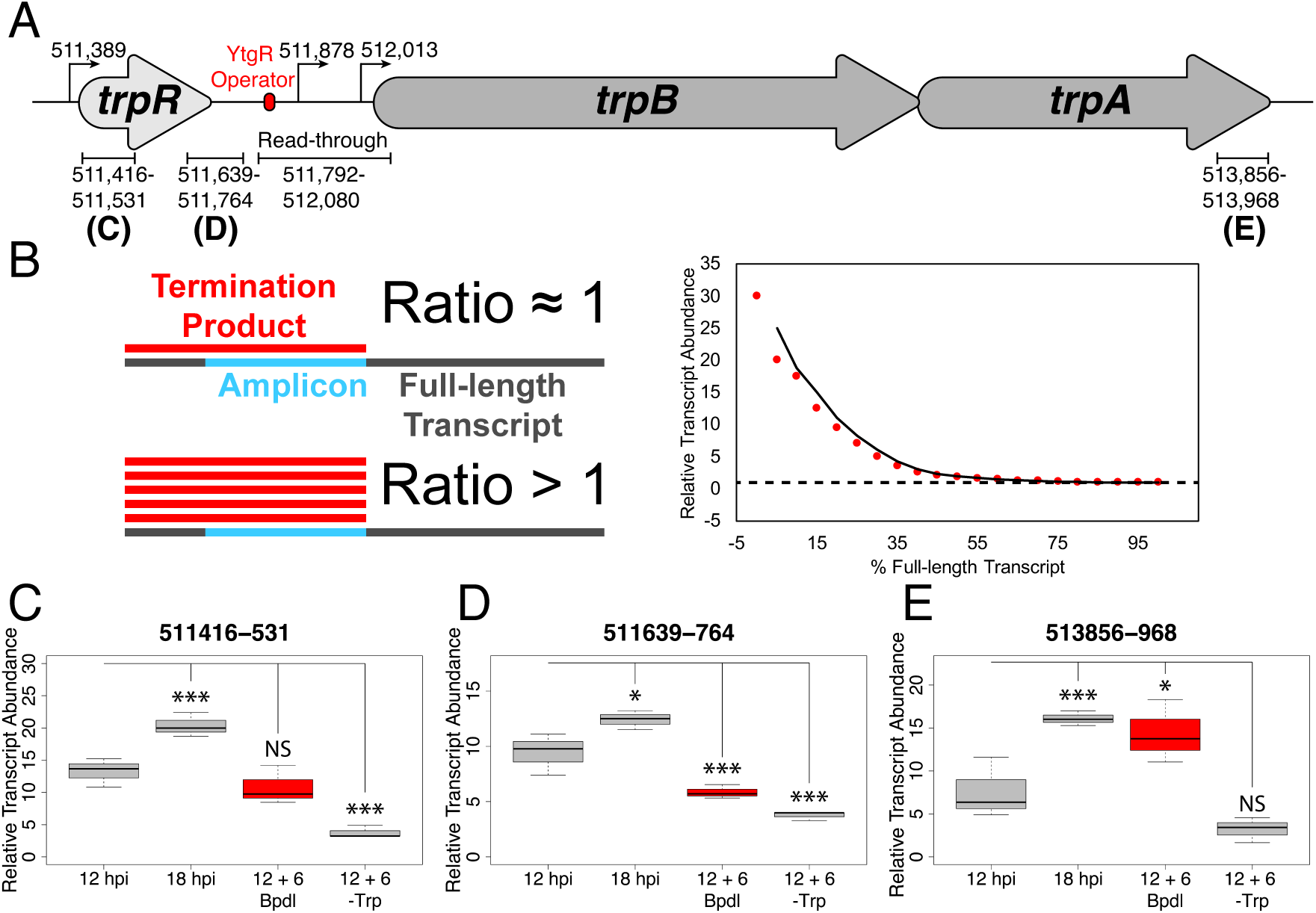
Iron limitation promotes transcription read-through at the YtgR operator site. (*A*) Cartoon depiction of amplicons analyzed in these experiments across the *trpRBA* operon, with the identified TSSs and YtgR operator site anootated. Drawn to scale. (*B*) Schematic representation of RT-qPCR analysis. On the left, a biological interpretation of the ratio used to determine the relative transcript abundance is provided. When a termination product (red) is poorly represented, *e.g.* read-through is high, the ratio of specific amplicon (blue) to full-length transcript (grey) should be close to 1.0. When the termination product is abundant, *e.g.* read-through is low, then the ratio should exceed 1.0. On the right, a graphical demonstration of this concept is provided using mock data. The dotted line represents the theoretical asymptote at a value of 1.0. (*C*) Analysis of transcription read-through by RT-qPCR was performed on 3’-RACE total RNA at three distinct loci across the *trpRBA* operon representing upstream transcription initiation (511,416-531), (*D*) YtgR operator site termination (511,639-764) and (*E*) terminal *trpBA* transcription (513,856-968). Abundance of each amplicon was normalized to a region (Read-through) predicted to be transcribed only as a part of the full-length product based on 5’ and 3’-RACE data (511,792-512,080). Thus, the ratio of each amplicon to the normalization amplicon represents the proportion of that amplicon encoded as part of the full-length transcript. At the YtgR operator termination site, iron limitation reduces the ratio relative to 12 hpi, suggesting that transcription read-through increases at this site under this condition. Statistical significance determined by One-way ANOVA followed by post-hoc pairwise t-tests. * = *p* < 0.05, ** = *p* < 0.01, *** = *p* < 0.005.

We first analyzed an amplicon from nucleotide 511,416 – 531 to monitor the relative abundance of transcript species associated with transcription initiating at the *trpR* promoter (Figure 7C). We observed that the representation of this amplicon was not significantly altered following iron limitation relative to 12 hpi, suggesting that the depletion of iron was not affecting initiation of transcription at the *trpR* promoter. Interestingly, at 18 hpi, the representation ratio of this amplicon significantly shifted further away from 1.0 (*p* = 0.00358), indicating that at 18 hpi this amplicon is represented less as a component of read-through transcription relative to 12 hpi. As expected, under Trp-deplete conditions, the representation ratio shifted significantly closer to 1.0 (*p* = 0.00064), consistent with read-through transcription of the full-length *trpRBA* message.

We then preformed the same analysis on an amplicon from nucleotide 511,639 – 764, immediately upstream of the TTS at the YtgR operator site to monitor condition-dependent read-through at this site (Figure 7D). We again observed that at 18 hpi, the representation ratio was significantly increased (*p* = 0.01046), and following Trp-depletion, the ratio was significantly decreased (*p* = 0.00023), as expected. Notably, and consistent with our hypothesis, we observed that the representation ratio of this amplicon was also significantly closer to 1.0 following iron limitation (*p* = 0.00407), suggesting that transcription read-through was increased at this site under iron limited conditions. Indeed, if YtgR is dissociating from the operator site during iron depletion, a greater proportion of transcripts would be expected to read-through this locus.

Finally, we analyzed an amplicon from nucleotide 513,856 – 968, at the very 3’- end of *trpA* to asses changes in terminal transcription under our experimental conditions (Figure 7E). At 18 hpi, we observed a significant increase in the representation ratio of this amplicon (*p* = 0.00476), which is likely attributable to both basal levels of alternative transcription from the IGR as well as poor transcription read-through of the full-length message. Following 6 hrs of Bpdl treatment, we also observed a significant increase in the representation ratio of this amplicon (*p* = 0.01510), which supports the finding that *trpBA* is being preferentially transcribed under this condition, distinct from the full-length *trpRBA* transcript. We were only able to detect a marginal decrease in the representation of this amplicon under Trp-depleted conditions (*p* = 0.07942), which may suggest that the very 3’-end of *trpRBA* is under-represented relative to our normalization amplicon, which falls within the middle of the operon. In fact, recent work has reported on the relatively poor representation of 3’-end mRNAs in *Chlamydia* (Ouellette, Rueden, & Rucks, 2016). In sum, this set of experiments provides evidence that iron-depletion specifically alters the representation of particular mRNA species across the *trpRBA* operon. Additionally, they implicate iron-dependent YtgR DNA-binding as the mediator of these effects. By alleviating YtgR repression via iron depletion, transcription is allowed to proceed through the operator site, albeit at basal levels. Concomitantly, transcription is specifically activated at the downstream alt. TSS for *trpBA.*

## Discussion

In this study, we provide a mechanistic explanation for the specific iron-limited induction of *trpBA* expression mediated by the repressor YtgR, representing a novel instance of integrated stress adaptation in *Chlamydia*. Utilizing an infected-epithelial cell culture model, we identified a previously undescribed iron-regulated promoter element within the *trpRBA* IGR responsible for the iron-limited induction of *trpBA* expression independent of *trpR.* Using *in silico,* biochemical and chemical-genetic methods, we demonstrate that YtgR binds the *trpRBA* IGR to regulate iron-dependent *trpBA* expression. Importantly, transcriptional repression in our heterologous system was shown to be dependent on an unaltered operator sequence that bears significant homology to the previously defined operator element in the *ytg* promoter. Previously published reports have demonstrated that YtgR is capable of directly binding DNA sequences containing this operator *in vitro* (Akers et al., 2011; Thompson et al., 2012). Furthermore, our infected-cell culture studies revealed that transcripts originating from the primary *trpR* promoter terminate within the IGR, notably at the putative YtgR operator site, and that transcription read-through at this locus is iron-dependent. Thus, we propose that YtgR regulates *trpBA* expression at two levels: repression of the *trpBA* promoter and premature termination of the major transcript generated from the *trpR* promoter (Figure 8). This is the first time an iron-dependent mode of regulation has been shown to control the expression of tryptophan biosynthesis in prokaryotes, which reflects the uniquely specialized nature of *C. trachomatis.*

**Figure 8.**
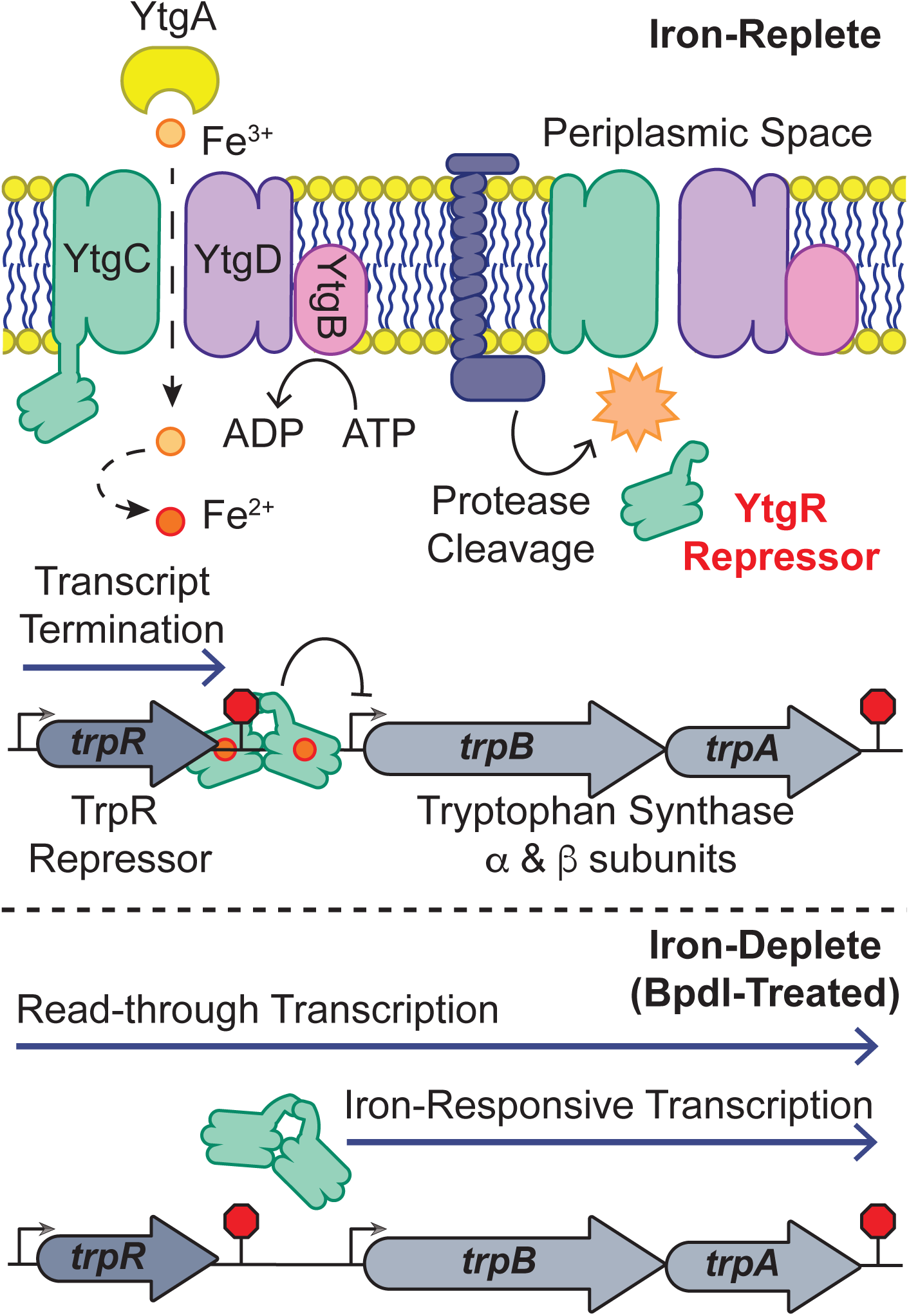
Model for proposed mechanism of iron-dependent YtgR-mediated regulation of *trpRBA* expression. Iron is imported through the YtgABCD ABC-type metal permease complex. YtgR is cleaved from the YtgCR permease-repressor fusion protein. In the presence of sufficient iron, holo-YtgR can bind to the *trpRBA* IGR to both terminate basal transcription from the primary *trpR* promoter and repress transcription initiation at the alternative *trpBA* promoter. Iron depletion inactivates YtgR DNA-binding, thus promoting read-through of basal transcription from the *trpR* promoter while also inducing transcription at the downstream *trpBA* promoter.

While we demonstrate here that iron-dependent *trpBA* expression originates from a novel promoter element immediately upstream of the *trpB* CDS, this is not the first description of an alt. TSS within the *trpRBA* IGR. Carlson, *et al.* identified an alt. TSS within the IGR which they suggested was responsible for *trpBA* expression (Carlson et al., 2006). In these studies, we were unable to confirm the presence of the previously identified alt. TSS by 5’-RACE. This is likely because Carlson, *et al.* examined the presence of transcript origins following 24 hrs of Trp starvation whereas here we monitored immediate responses to stress following only 6 hrs of treatment. Prolonged Trp depletion would result in a more homogeneously stressed population of chlamydial organisms that may exhibit the same preferential utilization of the promoter identified by Carlson, *et al.,* the detection of which is precluded in a more heterogeneous, transiently-stressed population. This may explain the observation of multiple T(S/T)Ss across the *trpRBA* operon in our studies. However, the contribution of such a Trp-dependent alt. TSS as identified by Carlson *et al.* to the general stress response of *C. trachomatis* remains unclear given its association with presumably abnormal organisms. Does utilization of this alt. TSS indicate abnormal growth or a *bona fide* stress adaptation? Moreover, Akers & Tan were unable to verify TrpR binding to the *trpRBA* IGR by EMSA, suggesting that some other Trp-dependent mechanism may control transcription from this site (Akers & Tan, 2006). Ultimately, our approach of investigating more immediate responses to stress revealed previously unreported mechanisms functioning to regulate Trp biosynthesis in *C. trachomatis*, underscoring the value of transient as opposed to sustained induction of stress.

Another mechanism of regulation reported to control the chlamydial *trpRBA* operon is Trp-dependent transcription attenuation. Based on sequence analysis, a leader peptide has been annotated within the *trpRBA* IGR (Merino & Yanofsky, 2005). Presumably, this functions analogously to the attenuator in the *E. coli trpEDCBA* operon; Trp starvation causes ribosome stalling at sites of enriched Trp codons such that specific RNA secondary structures form to facilitate RNAP read-thru of downstream sequences – in this case, *trpBA* (Yanofsky, 1981). However, robust experimental evidence to support the existence of attenuation in *C. trachomatis* is lacking. To date, the only experimental evidence that supports this model was reported by Carlson, *et al.,* who demonstrated that in a TrpR-mutant genetic background, an additional increase in *trpBA* expression could be observed following 24 hr Trp-depletion. (Carlson et al.,2006). However, this could be attributable to an alternative Trp-dependent, *but TrpR-independent* mechanism controlling *trpBA* expression at the alt. TSS identified by Carlson, *et al.* None of the data presented here point conclusively to the existence of a Trp-dependent attenuator. The additional termination sites identified in our 3’-RACE assay may represent termination events mediated by an attenuator, but without more specific analysis utilizing mutated sequences we cannot attribute attenuator function to those termination sites.

Interestingly, in *Bacillus subtilis*, Trp-dependent attenuation of transcription takes on a form markedly different from that in *E. coli.* Whereas attenuation functions in *cis* for the *E. coli trp* operon, *B. subtilis* utilize a multimeric <u>T</u>ryptophan-activated <u>R</u>NA-binding <u>A</u>ttenuation <u>P</u>rotein, TRAP, which functions in *trans* to bind *trp* operon RNA under Trp-replete conditions, promoting transcription termination and inhibiting translation (Gollnick, Babitzke, Antson, & Yanofsky, 2005). This interaction is antagonized by antiTRAP in the absence of charged tRNA^Trp^, leading to increased expression of TRAP regulated genes. We suggest that YtgR may represent the first instance of a separate and distinct clade of attenuation mechanisms: iron-dependent trans-attenuation. This mechanism may function independently of specific RNA secondary structure, relying instead on steric blockage of RNAP processivity, but ultimately producing a similar result. Possible regulation of translation remains to be explored. The recent development of new genetic tools to alter chromosomal sequences and generate conditional knockouts in *C. trachomatis* should enable a more detailed analysis of *trpRBA* regulation, including possible *trans*-attenuation (Mueller, Wolf, & Fields, 2016; Ouellette, 2018).

As a Trp auxotroph, what might be the biological significance of iron-dependent YtgR regulation of the *trpRBA* operon in *C. trachomatis*? We have already noted the possibility that iron-dependent *trpBA* regulation in *C. trachomatis* may enable the induction of a similar response to both Trp and iron starvation, stimuli likely mediated by IFN-γ *in vivo.* This mechanism also presents the opportunity for *C. trachomatis* to respond similarly to distinct *sequential* stresses, where a particular stress may prime the pathogen to better cope with subsequent stresses. To reach the female upper genital tract (UGT), where most significant pathology is identified following infection with *C. trachomatis,* the pathogen must first navigate the lower genital tract (LGT). *Chlamydia* infections of the female LGT are associated with bacterial vaginosis (BV), which is characterized by obligate and facultative anaerobe colonization, some of which produce indole (Sasaki-Imamura, Yoshida, Suwabe, Yoshimura, & Kato, 2011; Ziklo, Huston, Hocking, & Timms, 2016). This provides *C. trachomatis* with the necessary substrate to salvage tryptophan via TrpBA. Interestingly, the LGT is also likely an iron-limited environment. Pathogen colonization and BV both increase the concentration of mucosal lactoferrin (Lf), an iron-binding glycoprotein, which can starve pathogens for iron (Spear et al., 2011; Valenti et al., 2018). Lf expression is additionally hormone-regulated, and thus the LGT may normally experience periods of iron limitation (Cohen, Britigan, French, & Bean, 1987; Kelver et al., 1996). Moreover, the expression of TfR is constrained to the basal cells of the LGT stratified squamous epithelium (Lloyd, O’Dowd, Driver, & Tee, 1984), which likely restricts necessary Tf-bound iron from *C. trachomatis* infecting the accessible upper layers of the stratified epithelia (Nogueira, Braun, & Carabeo, 2017; Ouellette & Carabeo, 2010).

For *C. trachomatis,* iron limitation may therefore serve as a critical signal in the LGT, inducing the expression of *trpBA* such that Trp is stockpiled from available indole, allowing the pathogen to counteract impending IFN-g-mediated Trp starvation. We suggest the possibility that iron limitation in the LGT may be a significant predictor of successful pathogen colonization in the UGT. Unfortunately, testing these hypotheses in cell culture models of infection presents a significant challenge. Evaluating rescue of chlamydial growth in the presence of indole to specifically assess the iron-dependent role of *trpBA* requires simultaneous Trp and iron depletion. The former ensures indole utilization by the bacteria, and the latter de-represses YtgR-regulated *trpBA* expression. In theory, this is feasible, but in practice the combined stress rapidly induces aberrant development, muddying results obtained from such studies (data not shown). Ideally, genetic approaches could be employed to distinguish the regulatory effects of YtgR independent of TrpR. However, the genetic manipulation of trans-acting factors *(e.g.* YtgR) will presumably have unpredictable off-target effects. Genetically altering *cis-* acting factors - such as operator sequences - is more feasible, but at present we lack the information necessary to rationally mutate these sequences in *C. trachomatis* to interrogate these questions. The tight regulatory coordination at both the transcription initiation and termination steps would likely mean any mutation in the cis-acting sequences would affect both processes indiscriminately. Furthermore, *in vivo* infection models present challenges: attempting to answer these questions will likely require the use of *in vivo* non-human primate studies, as mouse models of *Chlamydia*-infection do not recapitulate immune-mediated Trp starvation (Nelson et al., 2005). Ultimately, these limitations do not undermine the biological significance of an iron-dependent mode of regulating Trp salvage, given the critical role played by this pathway during infection.

Finally, and of note, the expression of the ribonucleotide diphosphate-reductase-encoding *nrdAB* was also recently shown to be iron-regulated in *C. trachomatis* (Brinkworth et al., 2018). The regulation of *nrdAB* is known to be mediated by the presumably deoxyribonucleotide-dependent transcriptional repressor NrdR, encoded distal to the *nrdAB* locus (Case, Akers, & Tan, 2011). As NrdR activity is not known to be modulated by iron availability, this raises the intriguing possibility that here too a unique iron-dependent mechanism of regulation may integrate the chlamydial stress response to promote a unified response across various stress conditions. Future studies may require more metabolomics-based approaches to thoroughly dissect the integration of these stress responses, as transcriptome analyses alone often miss broader, pathway-oriented metabolic coordination. Ultimately, these studies point towards a need to carefully re-evaluate the molecular stress response in *Chlamydia,* with greater emphasis on the use of targeted approaches and treatment protocols that induce stress, but not persistence. We anticipate that the rapid progress of the field in recent years will continue to catalyze exciting and important discoveries regarding the fundamental biology of *Chlamydia.*

## Materials and Methods

### Eukaryotic Cell Culture and Chlamydial Infections

Human cervical epithelial adenocarcinoma HeLa (ATCC^®^ CCL-2) cells were cultured at 37° C with 5% atmospheric CO_2_ in Dulbecco’s Modified Eagle Medium (DMEM) supplemented with 10 μg/mL gentamicin, 2 mM L-glutamine, and 10% (*v/v*) filter sterilized fetal bovine serum (FBS). For all experiments, HeLa cells were cultured between passage numbers 4 and 16. *Chlamydia trachomatis* serovar L2 (434/Bu) was originally obtained from Dr. Ted Hackstadt (Rocky Mountain National Laboratory, NIAID). Chlamydial EBs were isolated from infected HeLa cells at 36-40 hours post-infection (hpi) and purified by density gradient centrifugation essentially as described (Caldwell, Kromhout, & Schachter, 1981).

For the infection of 6-well tissue culture plates, HeLa cells cultured to 80-90% confluency were first washed with pre-warmed Hanks Buffered Saline Solution (HBSS) prior to the monolayer being overlaid with inoculum (un-supplemented DMEM) at the indicated multiplicity of infection (MOI). Tissue culture plates were then centrifuged at 4° C with a speed of 1000 RPM (Eppendorf 5810 R table top centrifuge, A-4-81 rotor) for 5 minutes to synchronize the infection. Inoculum was aspirated and cells were washed again with pre-warmed HBSS prior to the media being replaced with pre-warmed complete DMEM. Infected cultures were then returned to the tissue culture incubator until the indicated times post-infection. This procedure was replicated exactly for the infection of 24-well tissue culture plates.

### Iron Starvation

*Chlamydia trachomatis* L2-infected HeLa cell cultures were starved for iron by supplementation of the media with the iron chelator 2,2-bipyridyl (Bpdl; Sigma Aldrich, St. Louis, MO, USA; CAS: 366-18-7) essentially as described (Thompson & Carabeo, 2011). Briefly, at the indicated times post-infection, infected cell cultures were washed with pre-warmed HBSS prior to the addition of complete DMEM (mock) or complete DMEM supplemented with 100 μM Bpdl. Infected cell cultures were returned to the incubator for the indicated treatment periods. Bpdl was prepared as a 100 mM stock solution in 100% ethanol and stored at -20° C for no longer than 6 months.

### Tryptophan Starvation

*Chlamydia trachomatis* L2-infected HeLa cell cultures were starved for tryptophan by replacement of complete DMEM with tryptophan-depleted medium. In brief, Tryptophan-replete or -deplete DmEM-F12 (U.S. Biological Life Sciences, Salem, MA, USA) powder media was prepared following manufacture instructions and supplemented with 10% (v/v) filter-sterilized FBS which had been previously dialyzed 16-20h at 4° C in PBS in a 10 kDa MWCO dialysis cassette. Media was then further supplemented with 10 μg/mL gentamicin. At the indicated times post-infection, complete DMEM was aspirated and wells were washed with pre-warmed HBSS prior to the addition of tryptophan-replete or –deplete medium. Infected cell cultures were returned to the incubator for the indicated treatment periods.

### Cloning

All constructs were cloned using standard molecular cloning techniques, *e.g.* restriction enzyme, homology-directed, etc. All primers and plasmids used in this study can be found in Supplementary file 3 and 4, respectively.

### Immunofluorescent Confocal Microscopy

At the indicated times post-infection, *C. trachomatis* L2-infected HeLa cell cultures seeded on glass coverslips in 24-well tissue cultures plates were first washed with prewarmed HBSS prior to fixation with 4% paraformaldehyde (PFA) in phosphate buffered saline (PBS) for 20 minutes at RT ° C. Fixation solution was aspirated and wells were washed with PBS prior to permeabilization with 0.2% Triton X-100 in PBS for 5 minutes at RT° C. Permeabilization solution was then decanted and cells were washed with PBS. The coverslips were blocked for 30 minutes with 1% bovine serum albumin (BSA) in PBS at RT° C. To stain for *Chlamydia*, coverslips were washed with PBS and PBS supplemented with 1% BSA and 1:500 convalescent human sera was added to wells and incubated at RT° C for 1 hour with rocking. Primary antibody solution was decanted and coverslips were again washed with PBS. Goat anti-human Alexa-647 (Invitrogen, ThermoFisher Scientific, Waltham, MA, USA) diluted 1:1000 in PBS with 1% BSA was then added to the wells and incubated in the dark for another hour at RT° C with rocking. Secondary antibody solution was then decanted, coverslips were washed again with PBS and coverslips were either immediately mounted on microscopy slides using ImmuMount (ThermoFisher Scientific) or VectaShield H-1000 (Vector Laboratories, Burlingame, CA, USA) or stored in the dark at 4° C until mounting. All images were acquired on a Leica TCS SP8 laser scanning confocal microscope, using identical settings, in the Integrative Physiology and Neuroscience Advanced Imaging Center at Washington State University. All images are Z-projections and were processed in Fiji (Schindelin et al., 2012) and Adobe Creative Suite identically for each comparative time-point.

### Nucleic Acid Preparation

RNA was harvested from *C. trachomatis*-infected HeLa cell monolayers by scraping 3 wells of a 6-well plate in ice-cold Trizol^®^ Reagent (ThermoFisher Scientific). Samples were then pooled and split into two technical replicates (RT-qPCR) or kept as one biological replicate (RACE). Trizol^®^-extracted samples were then thoroughly vortexed with a 100 μL volume of Zirconia beads prior to chloroform extraction. 100% ethanol was added to the aqueous phase and RNA was isolated using the Ambion^®^ RiboPure™ RNA Purification kit for bacteria following manufacturer instructions (ThermoFisher Scientific). DNA was removed from RNA samples using the Invitrogen DNA-*free*™ DNA Removal Kit following manufacturer instructions (ThermoFisher Scientific). RNA was stored at -20° C until further use.

cDNA was generated using either SuperScript^®^ IV Reverse Transcriptase (RT-qPCR; ThermoFisher Scientific) or SMARTScribe™ Reverse Transcriptase (RACE and RACE-specific qRT-PCR); Takara Bio, Kusatsu, Shiga Prefecture, Japan) essentially as described by the respective manufacturers. For cDNA generated for RT-qPCR, 650 ng of total RNA was used as a template in a 20 μL total reaction volume. For every RT reaction, a “no-RT” control, generated from 350 ng of total RNA template in a 10 μL total volume, was included. For 5’-RACE, cDNA was generated from 250 ng of total RNA using random primers in a 10 μL total volume and further processed in the RACE workflow. cDNA was stored at -20° C.

gDNA was harvested from *C. trachomatis*-infected HeLa cell monolayers by scraping 3 wells of a 6-well plate in ice-cold PBS + 10% Proteinase K (ThermoFisher Scientific). Samples were then pooled and split into two technical replicates for analysis of genome copy number by qPCR. gDNA was isolated using the DNeasy Blood and Tissue Kit following manufacturer protocols (QIAGEN, Hilden, Germany). gDNA was stored at -20° C until further use.

### Reverse Transcription Quantitative Polymerase Chain Reaction (RT-qPCR)

cDNA (or gDNA in qPCR), prepared as described above, was diluted 1:10 or 1:100 in nuclease-free H_2_O depending on the experimental condition being assayed *(e.g.* treatment, point in development cycle, etc.). On ice, 3.3 μL of diluted sample was added to 79 μL of PowerUp™ SYBR^®^ Green Master Mix (ThermoFisher Scientific) with specific qPCR primers diluted to 500 nM. From this master mix, each experimental sample was assayed in triplicate 25 μL reactions. Assays were run on an Applied Biosystems 7300 Real Time PCR System with cycling conditions as follows: Stage 1: 50.0° C for 2 min, 1 rep. Stage 2: 95.0° C for 10 min, 1 rep. Stage 3: 95.0° C for 15 sec, 40 reps. Stage 4: 60.0° C for 1 min, 1 rep. Primers were subjected to dissociation curve analysis to ensure that a single product was generated. For each primer set, a standard curve was generated using purified *C. trachomatis* L2 gDNA from EB preparations diluted from 2 x 10^-3^ to 2 x10^0^ ng per reaction. Ct values generated from each experimental reaction were then fit to standard curves (satisfying an efficiency of 95±5%) for the respective primer pair and from the calculated ng quantities, transcript or genome copy number was calculated as follows:

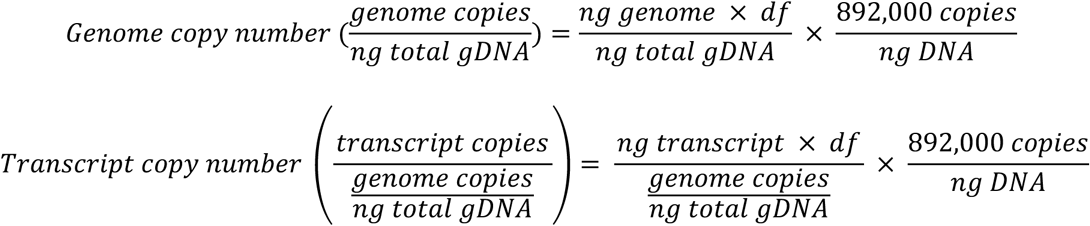

Where *df* = dilution factor and the number of copies/ng DNA is calculated based on the size of the *C. trachomatis* L2 genome assuming that the molar mass per base pair is 650 (g/mol)/bp (note that this value should be the same for any single-copy ORF on the genome). All quantifications of genome copy number were determined using the *ahpC* qPCR primer set. Values from replicate assays were averaged, and values from replicate RNA/gDNA isolations were averaged to obtain the mean and standard deviation for one biological replicate. For some experiments, to account for batch effects across biological replicates, data was transformed such that the mean of all samples in each replicate was identical. In some instances, batch correcting generated negative values, and in this case data sets were scaled such that the lowest value equaled 1.0. HeLa cells were infected at an MOI of 2 for all RT-qPCR studies.

### 5’ Rapid Amplification of cDNA Ends (5’-RACE)

All rAcE studies were performed using the SMARTer^®^ RACE 5’/3’ Kit (Takara Bio). To observe 5’-RACE products from the *trpRBA* operon, a “nested” RACE protocol was used as outlined in the SMARTer^®^ RACE 5’/3’ Kit user manual. Briefly, 1.25-2.5 μL of cDNA generated for RACE was added to a 25 μL reaction volume and run in a thermal cycler for 40 cycles using the touch-down PCR conditions described by the manufacturer. In brief, 5 cycles were run at an annealing temperature of both 72° C and 70° C prior to 30 cycles run with an annealing temperature of 68° C. Following this primary amplification, the RACE products were diluted 1:50 in Tricine-EDTA Buffer supplied by the manufacturer, and 2.5 μL of diluted primary RACE product was added to a 25 μL reaction volume and subjected to another 20 cycles of nested PCR, as described by the manufacturer, using primers designed within the amplicon of the primary RACE products. Samples were electrophoresed on a 2% agarose gel for visualization and analysis. HeLa cells were infected at a MOI of 5 for all RACE studies

### 3’ Rapid Amplification of cDNA Ends (3’-RACE)

3’-RACE studies were performed essentially identical to 5’-RACE with the exception that total RNA was subjected to poly(A) tailing with a Poly(A) Polymerase following manufacturer instructions (New England Biolabs, Ipswich, MA, USA). In brief, at least 3.5 μg of total RNA was incubated at 37° C with Poly(A) Polymerase in reaction buffer supplemented with ATP and murine RNase Inhibitor (New England Biolabs) for 30 minutes prior to heat-inactivation at 65° C for 20 minutes. RNA was re-isolated through an RNA clean-up filter cartridge (Ambion, ThermoFisher Scientific). A total of 125 ng of poly(A)-tailed total RNA was then used to generate 3’-RACE ready cDNA in a 10 μL reaction volume following manufacturer instructions. Primary and nested RACE was performed using 3’-RACE gene-specific primers following the same protocol for amplification described for 5’-RACE, with the exception that the extension time was adjusted to accommodate amplification of the full ∼3 kb *trpRBA* polycistronic message.

### Mapping of 5’/3’-RACE Products

5’-RACE products generated from either primary or nested RACE reactions were excised from the agarose gel and DNA was isolated using the NucleoSpin Gel and PCR Clean-up kit (Macherey-Nagel, Takara Bio). The isolated RACE products were then cloned into the pRACE vector supplied in the SMARTer^®^ RACE 5’/3’ Kit using the InFusion HD cloning kit (Takara Bio). Ligated vectors were transformed into chemically competent Stellar™ *E. coli* cells by heat shock. Transformed bacteria were plated on LB agar containing 50 μg/mL carbenicillin and incubated overnight at 37° C. Colonies were selected and screened for relevant inserts by PCR. Positive colonies were cultured overnight at 37° C in LB liquid broth containing 50 μg/mL carbenicillin and plasmids were isolated using the QIAprep Spin Miniprep kit (qIAGEN). Inserts were then sequenced by Eurofins Genomics using the default M13 Reverse sequencing primer. Returned sequencing data was aligned to the *C. trachomatis* L2 (434/Bu) genome (NCBI Accession: NC_010287) by BLAST and the most 5’ aligned nucleotide was considered the 5’ end of the insert. In the case of 3’-RACE data, the reverse complement sequence was first generated prior to alignment. Grouping of individual products was determined 1.) by clusters being greater than 30 nucleotides apart and 2.) by the specific RACE band that the alignment was derived from. These two criteria were not both satisfied in all cases and in those cases criteria 1.) was favored.

### Sequence Alignments

All *C. trachomatis* L2 434/Bu genome sequences were obtained from NCBI Accession NC_010287. Global pairwise sequence alignments were made using the EMBOSS Needle algorithm. Alignment parameters were set as follows: Matrix: DNAfull, Gap Open: 20, Gap Extend: 0.8, Output Format: pair, End Gap Penalty: True, End Gap Open: 10, End Gap Extend: 0.5. These conditions were sufficient to replicate the previously published alignment between the putative YtgR operator sequence and the TroR operator (Akers et al., 2011). Local pairwise sequence alignments were made using the EMBOSS Water algorithm. The putative YtgR operator was aligned to the entire 348 bp intergenic region of the *trpRBA* operon (C. *trachomatis* L2 [434/Bu] genome position 511,692-512,039). The alignment parameters were set as follows: Matrix: DNAfull, Gap Open:10, Gap Extend: 0.5, Output Format: pair. These are the default conditions and were chosen to remove bias from the alignment results.

### Two-Plasmid Reporter Assay

The YtgR-binding reporter assay was performed essentially as described, with minor modifications (Thompson et al., 2012). Promoter regions of interest were amplified from the *C. trachomatis* L2 (434/Bu) genome by PCR using the indicated primer sets, which included KpnI restriction endonuclease sites at the 5’ and 3’ ends of the promoter amplicon. The amplified fragments and the pCCT-EV plasmid were then KpnI-digested and the promoters ligated into the vector using T4 or Quick Ligase (New England BioLabs). Insert directionality was confirmed by directional colony PCR and positive clones were sequence verified. pCCT-*trpBA*ΔOperator was cloned by amplifying two fragments of the *pCCT-trpBA* vector with one ∼60mer primer containing the bases to be substituted for each fragment. Thus, the whole vector was split into two half-fragments containing the substituted bases. The two fragments were then cloned back together using In-Fusion Homology-Directed cloning (Takara Bio) to yield the final vector. Electrocompetent BL21(DE3) *E. coli* (Sigma Aldrich) were co-transformed by electroporation with the pCCT reporter plasmid and the pET151 expression vector (-EV or –YtgR) and plated on double selective LB agar containing 50 μg/mL carbenicillin and 15 μg/mL tetracycline. Prior to plating of transformed cells, 50 μL of 40 mg/mL X-Gal in DMSO (EMD Millipore, Burlington, MA, USA) was applied to the plate for colorimetric determination of β-galactosidase expression. Transformants were incubated overnight at 37° C. The following evening, blue colonies from each experimental condition were selected and cultured overnight in LB liquid broth containing 0.2% (*w/v*) D-glucose (for catabolite repression of expression vectors), 50 μg/mL carbenicillin and 15 μg/mL tetracycline. Cultures were incubated overnight at 37° C. The following morning, overnight cultures were spun down to remove glucose-containing media and subcultured in LB liquid broth medium containing 50 μM FeSO_4_, 50 μg/mL carbenicillin and 15 μg/mL tetracycline to an OD_600_ of 0.45. Cultures were incubated for 1 hour at 37° C and sub-cultured a second time in the same media to an OD_600_ of 0.1. Cultures were returned to the incubator for another hour prior to the addition of 500 μM isopropyl β-D- 1-thiogalactopyranoside (IPTG) to induce pET151 expression from the *lac* promoter. Cultures were incubated another hour prior to the addition of 0.2% L-arabinose to induce *lacZ* expression from the *araBAD* promoter. Cultures were incubated a final 2 hours prior to the collection of a 0.1 mL volume of cells for assaying β-galactosidase activity by the Miller Assay (J. H. Miller, 1972). Cell pellets were stored at -80° C prior to being assayed. To assay β-galactosidase activity, cell pellets were first re-suspended in Z-buffer (pH 7.0, 60 mM Na_2_HPO_4_, 40 mM NaH_2_PO_4_, 10 mM KCl, 1 mM MgSO_4_ and 2.7 μL/mL β-mercaptoethanol). 50 μL of 0.1% SDS and 100 μL of chloroform were then added to each sample prior to thorough vortexing. Samples were equilibrated for 5 minutes at 30° C and 200 μL of 4 mg/mL ortho-nitrophenyl-β-galactoside (ONPG) prepared in Phosphate Buffer (pH 7.0, 60 mM Na_2_HPO_4_, 40 mM NaH_2_PO_4_) were added to the samples to initiate the reaction. Reactions were stopped by the addition of 500 μL 1 M Na_2_CO_3_. Absorbance was measured on a FLUOStar Optima plate reader (BMG Labtech, Offenburg, Germany) at 420 nm and Miller Units were calculated as:

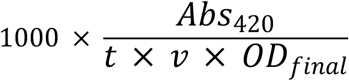

Where *t* = reaction time, *v* = volume of cells and *OD_final_* = OD_600_ at the time of sample collection. It was empirically determined that the subtraction of absorbance at 550 nm had a negligible effect on the calculated value. A blank sample lacking cells was included in each experimental batch and used as a reference for absorbance. For each experimental condition, three independent co-transformed colonies were assayed in technical triplicate. In some instances, significantly high Miller Unit outliers were excluded by Grubb’s Test (*p* < 0.05) under the assumption that extreme *lacZ* expression may reflect plasmid copy number or reporter gene expression issues.

### RNA-Sequencing

RNA-Sequencing experiments were performed as described in their original publication (Brinkworth et al., 2018). Coverage maps were generated by mapping all reads across three biological replicates to a single reference file in CLC Genomics Workbench v11. To facilitate easy analysis of IGR boundaries, the *C. trachomatis* L2 434/Bu genome (Accession: NC_010287) was modified to contain annotations for non-operonic intergenic regions, and this genome was used as the reference for read mapping. Read mapping was performed using default settings in CLC Genomics Workbench. Data aggregation in the Reads track was set to aggregate above 1bp.

### Graphs and Statistical Analysis

All graphs were generated using the ggplot2 package (Wickham, 2009) in R Studio, in Excel and/or in the Adobe Creative Suite. All line plots and bar graphs represent the mean ± one standard deviation unless otherwise noted. All box and whisker plots represent the distribution of data between the 1^st^ and 3^rd^ quartile range within the box, while the whiskers represent data within 1.5 interquartile ranges of the 1^st^ or 3^rd^ quartile. Extreme values outside this range are plotted as open circles. The 2^nd^ quartile (median) is plotted as a black line within the box. Histogram plots were generated with a bin width of 20 and are plotted on a density scale. The overlaid density plots represent a statistical approximation of the data over a continuous scale. All statistical analyses were carried out in R Studio. All statistical computations were performed on the mean values of independent biological replicates calculated from the indicated number of respective technical replicates. For single pairwise comparisons, a two-sided unpaired Student’s *t*-test with Welch’s correction for unequal variance was used to determine statistical significance. For multiple pairwise comparisons, a One-Way Analysis of Variance (ANOVA) was conducted to identify significant differences within groups. If a significant difference was detected, then the indicated post-hoc pairwise test was used to identify the location of specific statistical differences. A *p*-value less than 0.05 was considered statistically significant. For all figures, * = *p* < 0.05, ** = *p* < 0.01, and *** = *p* < 0.005.

## Acknowledgements

We thank Liam Caven, Korinn Murphy and Matthew Romero for critical review of this manuscript; Dr. Christopher C. Thompson for the establishment of the *E. coli* YtgR reporter system and generation of the pCCT construct; Dr. Scot P. Ouellette for support, critical feedback and advice; and Dr. David Dewitt for expert advice, training and maintenance of equipment in the IPN Advanced Imaging Center. This work was supported by NIH grants R01-AI065545 to R.A.C.; 1F31AI136295 and 5T32GM008336 to N.D.P.; N.D.P. was also supported by an Achievement Reward for College Scientists (ARCS; Seattle Chapter) Fellowship.

## Author Contributions

N.D.P. and R.A.C. wrote the manuscript; A.J.B. and R.A.C. designed and analyzed data for the RNA-Sequencing experiments; A.J.B. performed RNA-Seq experiments; N.D.P. and R.A.C. re-analyzed the RNA-Sequencing data for this publication; N.D.P. and R.A.C. designed all other experiments; N.D.P. performed all other experiments; N.D.P. and R.A.C. analyzed and interpreted the data for all other experiments.

## Supplementary Figures

**Figure 1 – Figure Supplement 1.**
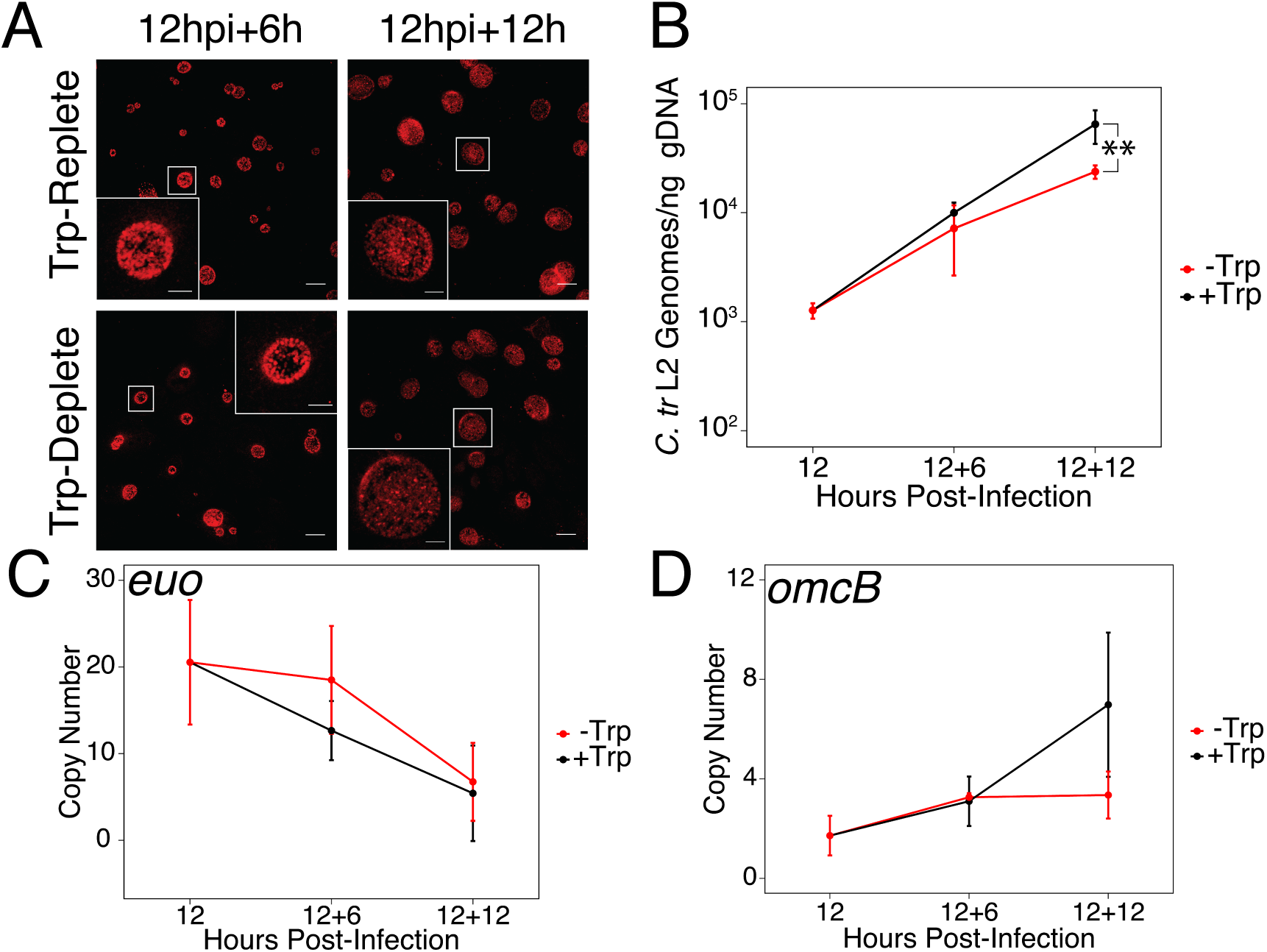
Brief media-defined tryptophan limitation does not produce characteristically persistent *Chlamydia. (*A*) C. trachomatis* L2-infected HeLa cells were fixed and stained with convalescent human sera to image inclusion morphology by confocal microscopy following tryptophan limitation at the indicated times post-infection. Figure shows representative experiment of two biological replicates. Scale bar = 15 μm, Inset scale bar = 5 μm. (*B*) Genomic DNA (gDNA) was harvested from infected HeLa cells at the indicated times post-infection under tryptophan-replete (black) and -deplete (red) conditions. Chlamydial genome copy number was quantified by qPCR. Chlamydial genome replication is stalled following 12 hours of tryptophan limitation, but not 6. N=2. (*C*) Total RNA was harvested from infected HeLa cells at the indicated times post-infection under tryptophan-replete (black) and -deplete (red) conditions. The transcript abundance of hallmark persistence genes *euo* and (*D*) *omcB* were quantified by RT-qPCR and normalized against genome copy number. No period of tryptophan limitation significantly impacted *euo* or *omcB* expression. N=2. Statistical significance was determined by One-Way ANOVA followed by post-hoc pairwise *t*-tests with Bonferroni’s correction for multiple comparisons.

**Figure 4 – Figure Supplement 1.**
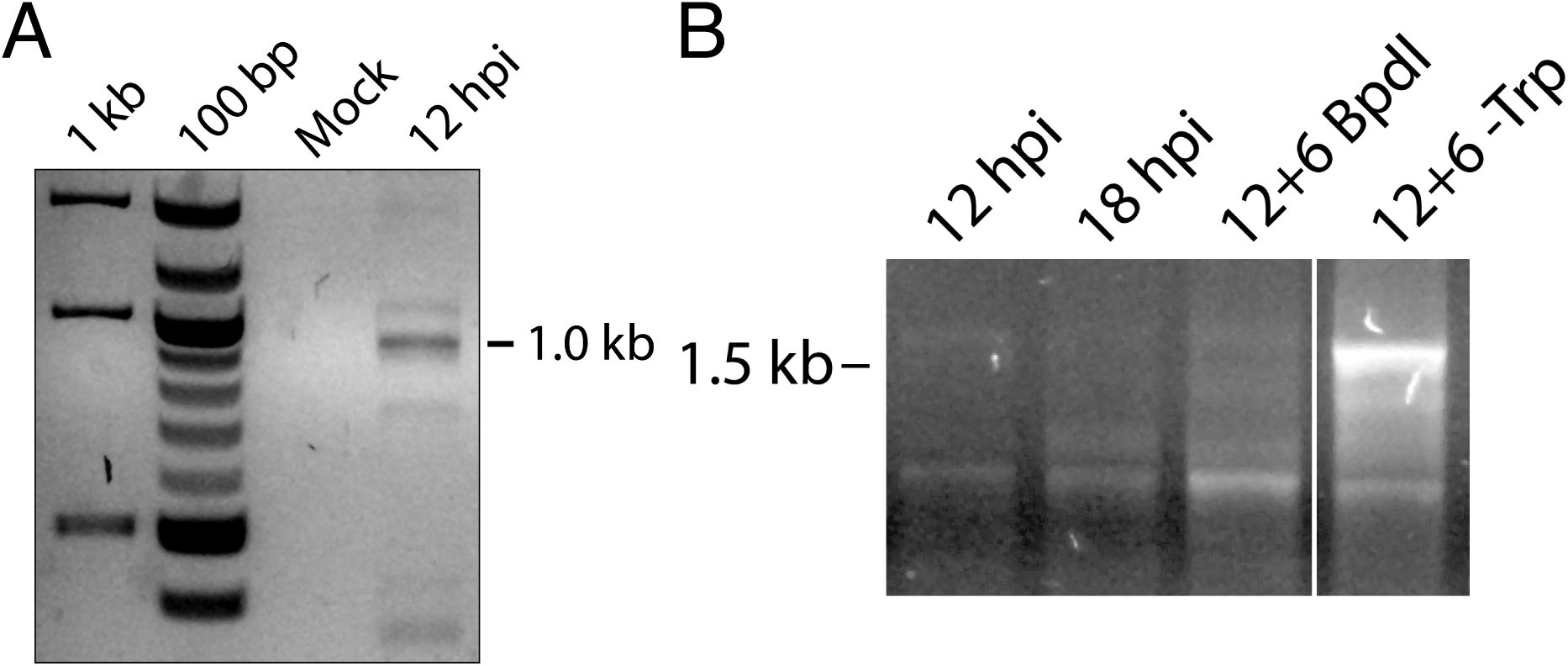
5’-RACE conditions produce *Chlamydia-specific* products that are amplified in primary RACE. (*A*) Total RNA harvested from mock-infected HeLa cells was processed for 5’-RACE in parallel with infected samples to determine specificity of amplified products to *Chlamydia*-infected cells. No RACE products were detected in the mock-infected sample. (*B*) Primary products amplified from 5’-RACE were electrophoresed on an agarose gel and visualized. Weak bands corresponding to those detected by nested 5’-RACE were observed, with the noted relative abundance of the 1.0 and 1.5 kb products in the Bpdl-treated and Trp-deplete conditions, respectively.

**Figure 4 – Figure Supplement 2.**
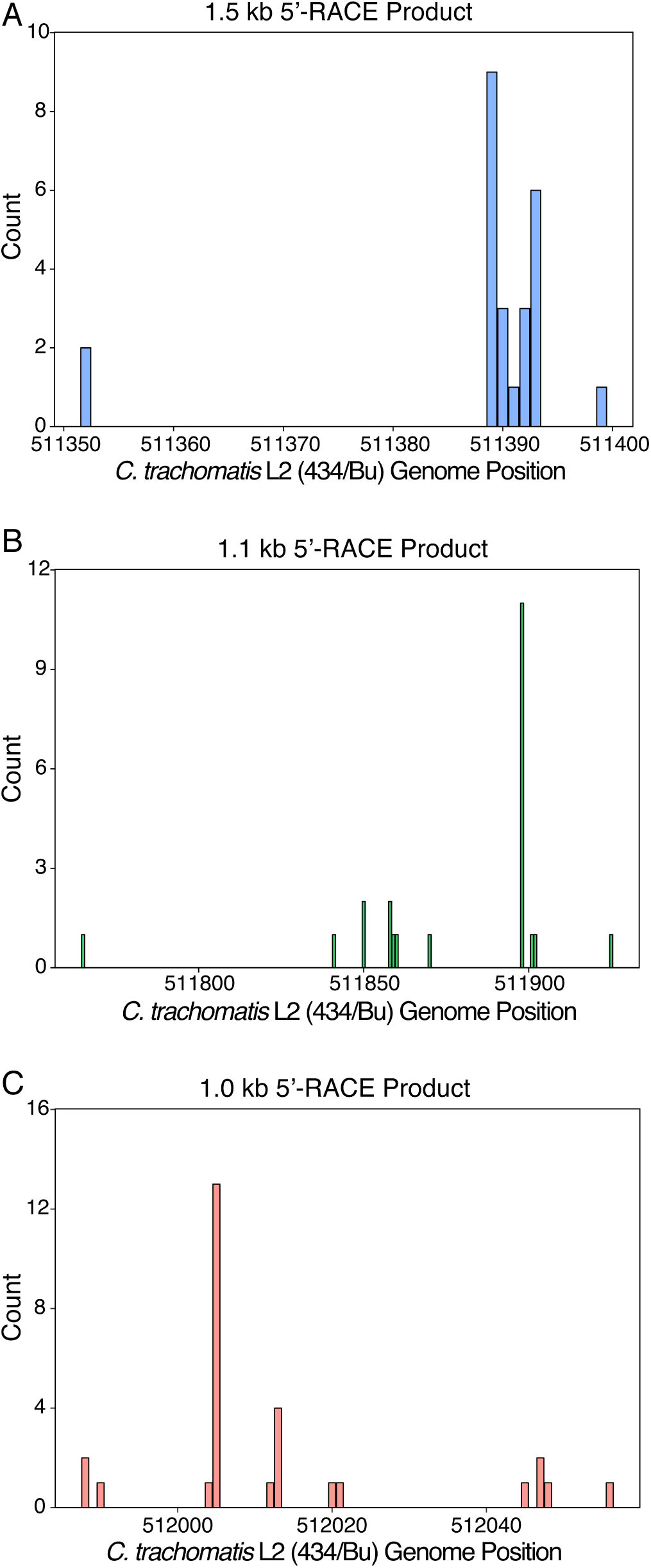
Mapping of 5’-RACE products at the individual nucleotide level. (*A*) Mapping of the 1.5 kb 5’-RACE product. (*B*) Mapping of the 1.1 kb 5’-RACE product. (*C*) Mapping of the 1.0 kb 5’-RACE product.

**Figure 5 – Figure Supplement 1.**
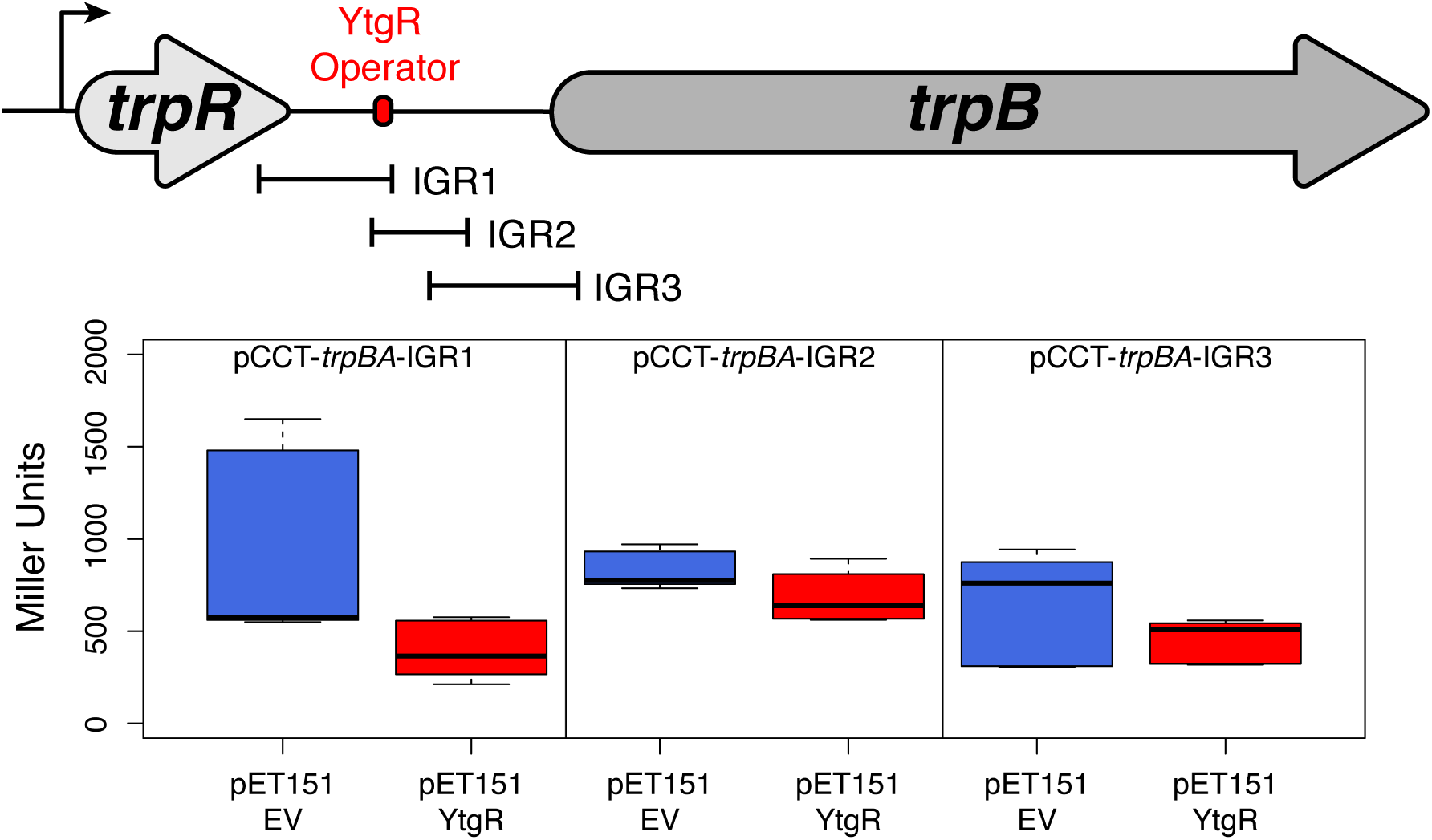
Truncated fragments of the *trpRBA* IGR are not sufficient to confer YtgR repression phenotype, regardless of the presence of the putative operator site. The cartoon depiction of the *trpRBA* operon indicates the position of the IGR fragments cloned into each of the reporter vectors *(i.e.* pCCT-IGR1 through IGR3). No statistically meaningful differences were detected in the β-galactosidase activity in the presence of any of the IGR fragments.

**Figure 6 – Figure Supplement 1.**
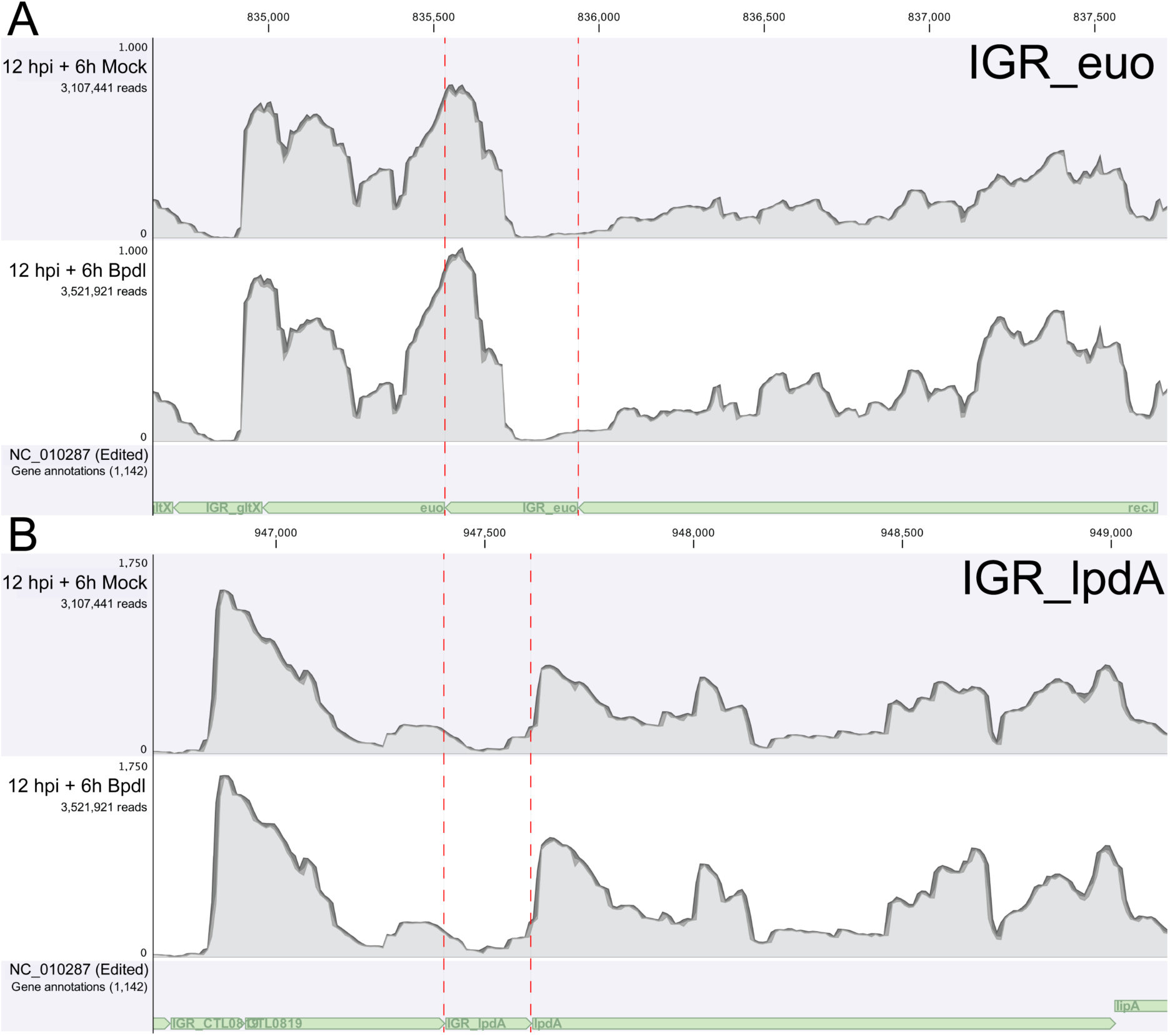
RNA-Sequencing coverage map of reads mapping to the intergenic regions upstream of *euo* and *lpdA*. (*A*) Coverage map for the IGR upstream of *euo*, a gene that is neither iron-regulated nor YtgR-regulated. (*B*) Coverage map for the IGR upstream of *lpdA*, a gene that is iron-regulated but not YtgR-regulated. Neither coverage map indicates an increase in reads mapping to these intergenic regions, suggesting that the effect observed at the *trpRBA* IGR is specific.

**Figure 6 – Figure Supplement 2.**
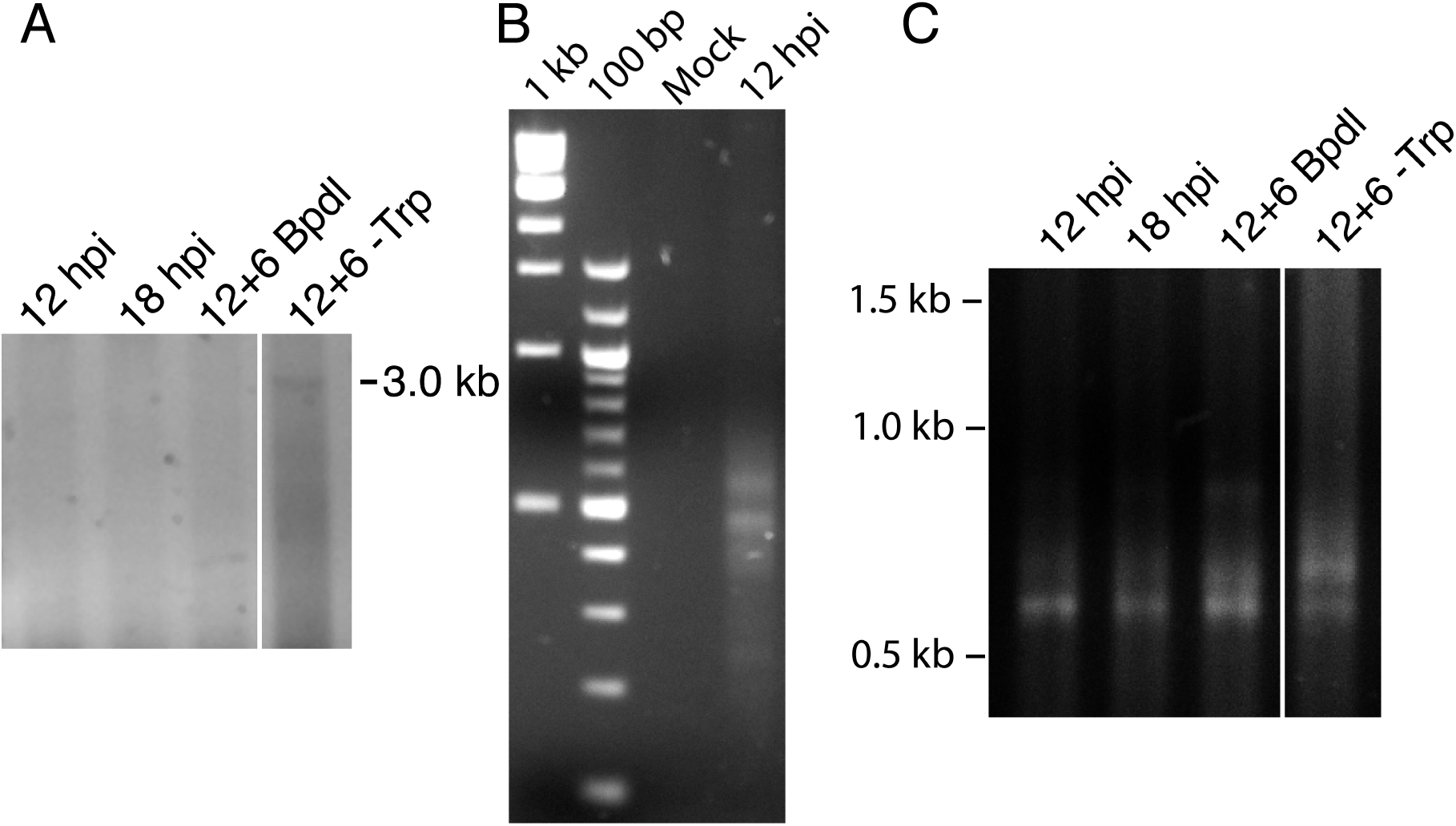
3’-RACE conditions produce *Chlamydia*-specific products that are amplified in primary RACE. (*A*) Contrast-adjusted image to show weak amplification of ∼3.0 kb full-length *trpRBA* transcript in nested 3’-RACE reaction. (*B*) Total RNA harvested from mock-infected HeLa cells was processed for 3’-RACE in parallel with infected samples to determine specificity of amplified products to *Chlamydia*-infected cells. No RACE products were detected in the mock-infected sample. (*C*) Primary products amplified from 3’-RACE were electrophoresed on an agarose gel and visualized. Weak bands that were relatively non-specific were detected in the 3’-RACE primary amplification, emphasizing the utility of producing enhanced specificity in the nested amplification.

**Figure 6 – Figure Supplement 3.**
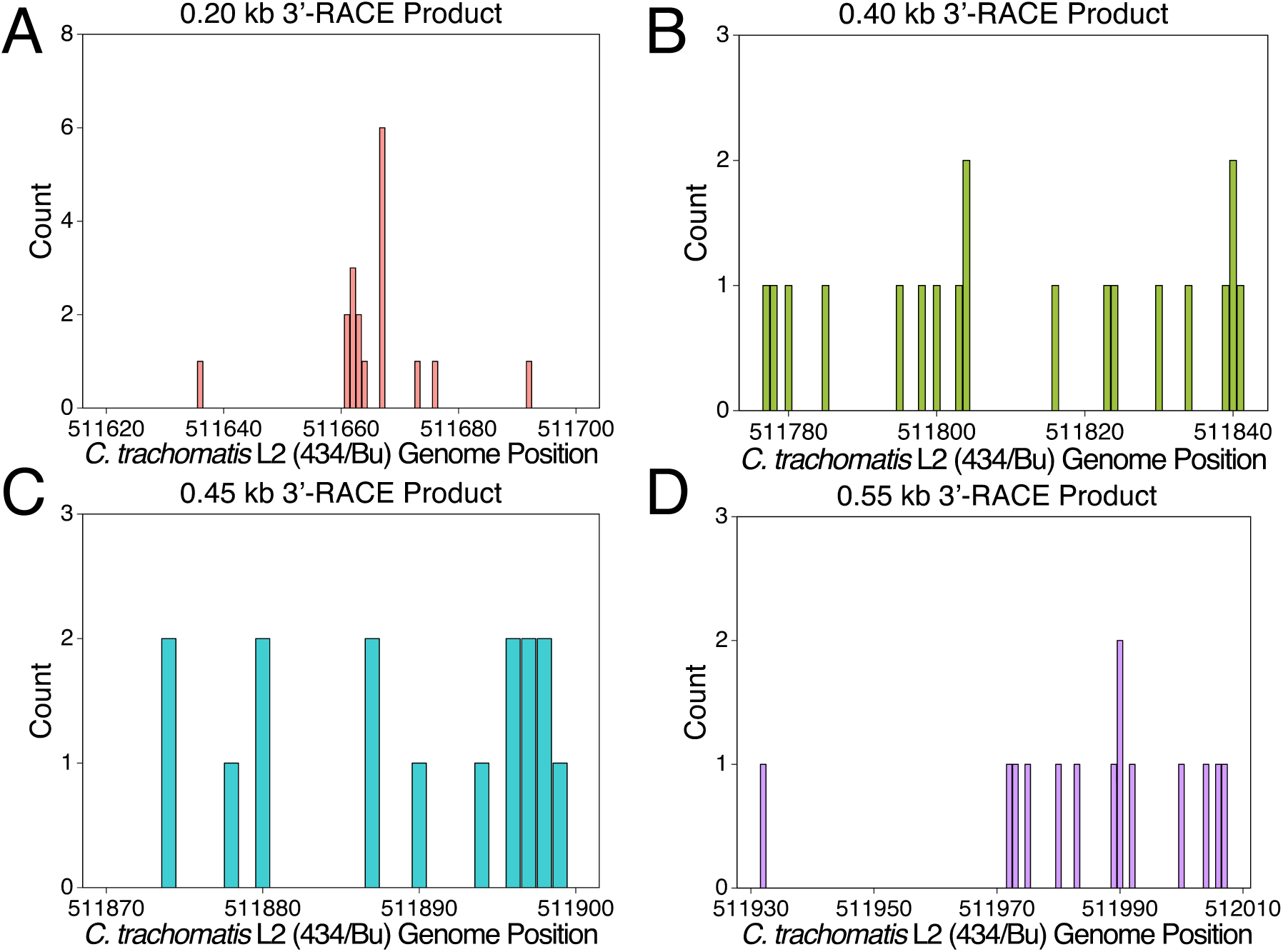
Mapping of the 3’-RACE products at the individual nucleotide level. (*A*) Mapping of the 0.20 kb 3’-RACE product. (*B*) Mapping of the 0.40 kb 3’-RACE product. (*C*) Mapping of the 0.45 kb 3’-RACE product. (*D*) Mapping of the 0.55 kb 3’-RACE product.

